# BlueBerry: Closed-loop wireless optogenetic manipulation in freely moving animals

**DOI:** 10.1101/2025.01.30.635697

**Authors:** Ali Nourizonoz, Benoit Girard, Maëlle Guyoton, Gregorio Galiñanes, Raphael Thurnherr, Sebastien Pellat, Camilla Bellone, Daniel Huber

**Affiliations:** University of Geneva, Department of Basic Neurosciences, Geneva, Switzerland

## Abstract

Optogenetics is a powerful approach for linking neural activity to behavior by enabling precise manipulation of defined neuronal populations. Recent advances in wireless technologies have led to remotely controlled optogenetic devices that facilitate the study of more complex behaviors, including social interactions. However, implementing real-time, automated, closed-loop experiments in large-scale, naturalistic environments and in groups of interacting animals remains challenging. Moreover, many existing devices are difficult to reproduce, requiring specialized engineering expertise or fabrication facilities that are not widely available. Here we introduce BlueBerry (www.OptoBlueBerry.org), a lightweight (1.4 g), battery powered, multi-channel wireless optogenetic device that is openly available for the scientific community and can be assembled entirely from off-the-shelf-components. BlueBerry combines robust long-range wireless communication with flexible control of stimulation parameters, making it particularly suited for behavior-triggered optogenetic experiments in large arenas and in multiple freely interacting animals. We demonstrate that BlueBerry can not only guide decision making during large-scale navigation, but also deliver individually controlled, behavior-triggered optogenetic stimulation to multiple freely interacting mice, enabling modulation of social dynamics in real-time. The open availability, modular design, and straightforward integration with existing behavioral frameworks make BlueBerry a practical and scalable tool for systems neuroscience.

## Introduction

Understanding the link between brain and behavior requires detailed investigation of the related neural circuits (Krakauer et al. 2017). Although neural recordings play a crucial role in observing correlates of behaviors, manipulating brain circuits is essential to establish the causality of involved neural networks. To this end, neuroscientists have widely adopted optogenetic techniques for precise and controlled manipulation of neuronal activity in behaving animals (Boyden et al. 2005; Deisseroth 2011).

In freely moving conditions, optogenetic stimulation was initially implemented using tethered light sources (Huber et al. 2008) or via optical fibers connected to external lasers (Adamantidis et al. 2007; Gradinaru et al. 2009; Tsai et al. 2009). The development of compact and power-efficient electronics enabled the development of integrated wireless optogenetic devices (Kim et al. 2013; Park et al. 2015; Montgomery et al. 2015; Jeong et al. 2015), reducing physical constraints on the animals and extending the range of behaviors to complex three-dimensional environments (S.-G. Park et al. 2018) or social settings (Yang et al. 2021; Li et al. 2022). In such contexts, traditional tethered setups are impractical, as cables restrict movements and can easily tangle with structures in naturalistic arenas or with other animals.

Wireless optogenetic devices can mainly be categorized in battery-free or battery powered designs. The primary benefit of using battery-free wireless optogenetics (Kim et al. 2013; Yang et al. 2021; Montgomery et al. 2015; Mickle et al. 2019; Shin et al. 2017) is their capability to operate for extended periods, making them ideal for experimental setups that require long experiment durations. However, their range is typically restricted to relatively small spaces as magnetic induction coil systems have to be installed around the experimental arena, limiting their use in large-scale or complex volumetric environments (Shin et al. 2017). Finally, these devices are often chronically implanted as a single integrated unit, which can be more invasive and less flexible, and may hinder reuse across animals (Yang et al. 2021).

In parallel, battery-powered wireless optogenetic systems have been developed for freely moving animals (Kathe et al. 2022; Li et al. 2022; Michoud et al. 2021). This approach generally offers a larger operational range, often extending to several meters, without the need for specialized power coils, and is not restricted to planar laboratory cages. Critically, only the miniature light sources need to be implanted; the battery and control electronics can remain detachable. This modularity facilitates the design of minimally invasive light-delivery systems targeting superficial or deep brain regions and allows the reuse of the same device across animals.

Despite these advances, several limitations prevent widespread adoption of wireless optogenetics. Many systems are difficult to reproduce because detailed build instructions are lacking, or the devices are not fully open or they depend on specialized equipment or cleanroom fabrication (McCall et al. 2013; Yang et al. 2022, Kathe et al. 2022; Michoud et al. 2021; Li et al. 2022; Jeong et al. 2015). Finally, many devices are tailored for specific tasks or limited to specific experimental arenas (Mickle et al. 2019; Kathe et al. 2022; Michoud et al. 2021), complicating their integration into diverse behavioral frameworks, particularly those requiring real-time, behavior-triggered stimulation in complex or large-scale environments, or in groups of interacting animals.

To address these gaps, we developed BlueBerry (www.OptoBlueBerry.org), a battery-powered multi-channel wireless optogenetic system controlled and programmed remotely through standard Bluetooth Low Energy (BLE) communication protocol. BlueBerry is small, lightweight, low-cost, and fully documented, and can be assembled using only commercially available off-the-shelf components. Stimulation patterns (e.g., frequency, pulse width, and pulse count) can be configured independently for each channel and device via a dedicated Arduino-based control unit, BlueHub, which interfaces with existing behavioral setups through simple TTL inputs.

We demonstrate BlueBerry’s capabilities in two closed-loop, behavior-triggered paradigms in freely moving mice. First, we show that optogenetic stimulation of the somatosensory cortex can guide navigation in large-scale “infinite” Y-mazes based on artificial sensory cues. Second, we use real-time tracking of multi-animal social interactions to reinforce specific agonistic behaviors by stimulating ventral tegmental area dopaminergic (VTA-DA) neurons, transiently reshaping social dynamics in groups of mice. Together, these experiments highlight BlueBerry as an accessible, versatile, and scalable tool for naturalistic, closed-loop optogenetic studies in freely moving animals.

## Results

### Implementation of BlueBerry wireless optogenetic system

To wirelessly deliver optogenetic stimulation to freely moving mice in large scale or multi-animal settings, we developed a compact (14 x 10.5 x 7 mm), lightweight (1.4 g), battery-powered device capable of controlling up to two independent channels (Fig. 1a). We named our device BlueBerry as it is the size and the weight of a blueberry and uses bluetooth for wireless communication.

**Figure 1.**
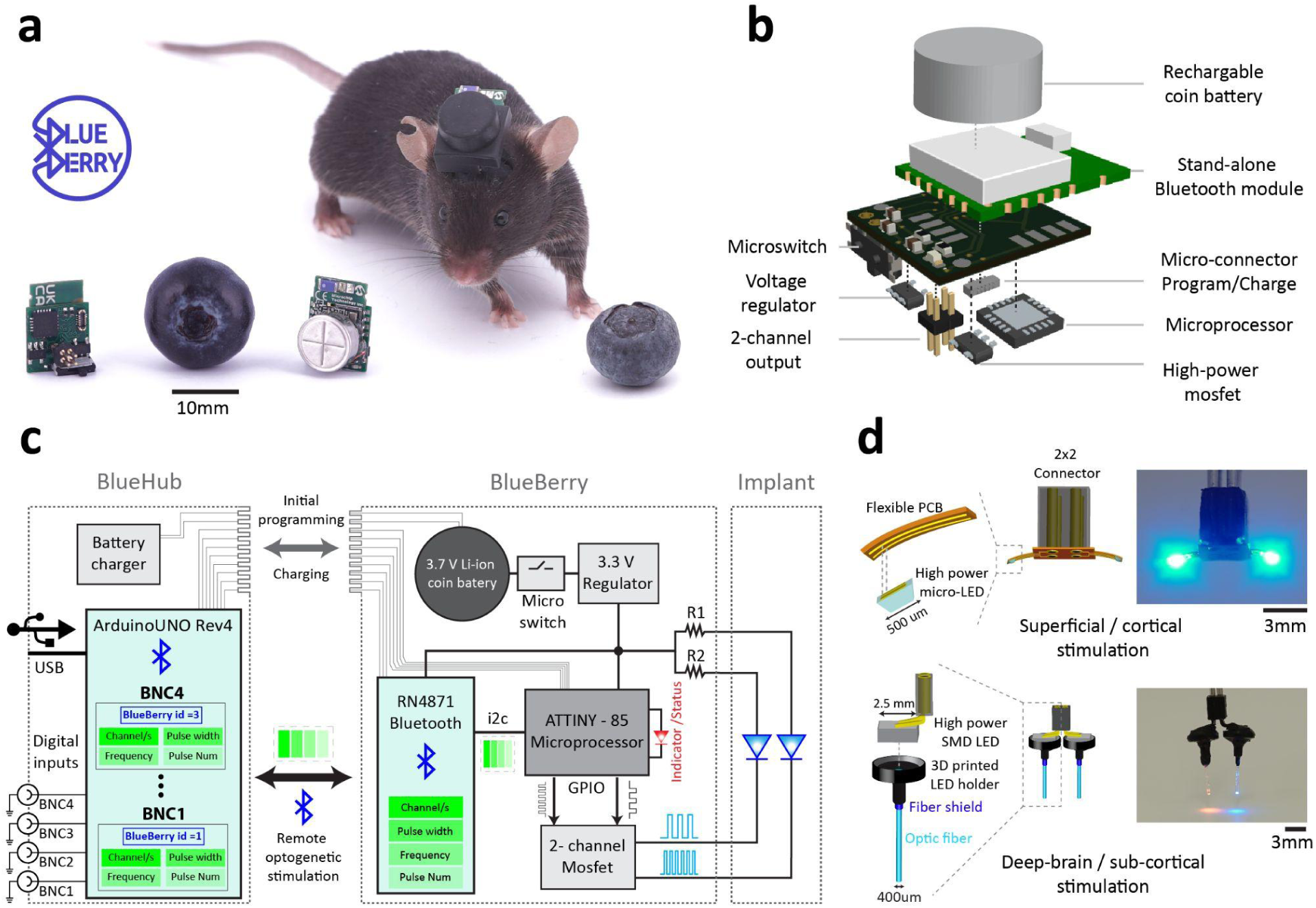
Description of BlueBerry wireless optogenetic system. **a)** Front and back of the BlueBerry device shown beside a real blueberry for scale (left). Image of a mouse carrying a BlueBerry device connected to the multichannel LED implant fixed on the skull (right). **b)** Exploded 3D view of the BlueBerry device illustrating all miniature electronic components. **c)** Simplified schematic of the control BlueHub control unit, the BlueBerry device and LED implants. Each BlueHub input channel (BNC1–4) can be assigned to a function that includes the BlueBerry (BlueBerry id) device to activate and the corresponding stimulation parameters (stimulation channel, frequency, pulse width and number of pulses). When a channel is triggered, the BlueHub sends a Bluetooth packet to the assigned BlueBerry device containing the specified stimulation protocol. On the BlueBerry side, Bluetooth packets are decoded inside the bluetooth module and communicated with the microcontroller where each output stimulation channel is configured in terms of frequency, pulse width and number of pulses. **d)** Multichannel implants targeting superficial cortical layers (top) through surface mount micro LEDs or deep brain regions through custom made fiber coupled LED (bottom).

The BlueBerry includes a dedicated stand-alone bluetooth module communicating with a microcontroller through a custom designed printed circuit board (PCB, Fig. 1b and Supplementary Fig. 1 & 2) to deliver high-power pulses to both output channels with latencies below 30 ms for distances up to 3 m (Supplementary Fig. 6). Stimulation protocols such as pulse-width and frequency (independently for each stimulation channel) can be configured on-the-fly for each BlueBerry device. Under typical operating conditions, a single charge supports up to 7 hours of use (Supplementary Fig. 6).

Programming and control are handled by the BlueHub, a user-friendly Arduino-based unit (Fig. 1c; Supplementary Fig. 3–4). The BlueHub features up to four digital input channels, each of which can be connected to an arbitrary external behavioral framework (3–5 V TTL). Each input can be mapped to a specific BlueBerry device and its corresponding stimulation protocol (Fig. 1c). When a given input channel is triggered, BlueHub sends a BLE packet containing the specified stimulation parameters to the assigned BlueBerry device. Onboard firmware on the BlueBerry decodes incoming packets and configures the two output channels accordingly.

BlueBerry connects via a standard 2 × 2 connector to custom LED-based implants (Fig. 1d). For superficial cortical stimulation, we used off-the-shelf micro-LEDs (500 x 500 μm, pulses up to 60 mW) mounted on flexible PCBs placed directly over the target region. For deep-brain stimulation, LEDs were coupled to a 400 µm optical fiber, achieving 7–9 mW at the fiber tip (Supplementary Fig. 5). Together, this modular architecture allows BlueBerry to be easily integrated into existing behavioral frameworks, with behavioral events triggering low-latency stimulation of either superficial or deep brain regions in freely moving animals.

### Closed-loop manipulation of large-scale navigation

Artificial sensory cues delivered via optogenetic stimulation to the sensory cortex provide interesting means to probe the limits of specific sensory pathways. We used BlueBerry to deliver such cues to primary somatosensory barrel cortex during large-scale navigation.

Mice (N = 6) expressing channelrhodopsin (ChR2) bilaterally in the barrel cortex were implanted with flexible two-channel micro-LED implants (Fig. 2a; Supplementary Fig. 5). Each micro-LED was positioned over the targeted cortical area (Fig. 2a–b). Inspired by the classic infinite virtual maze system for insects (Hassenstein and Reichardt 1956), we designed two elevated infinite Y-mazes of different sizes (1mx1m and 2.5 m x 2.5 m, Supplementary Fig. 7) consisting of arms containing a water port as reward at their center and connected through four intersections (Fig. 2c). In this paradigm, water-restricted mice arriving at any intersection triggered the activation of a directional optogenetic cue, by activating either the left or the right micro-LED (stimulating the contralateral sensory cortex, see Methods). Mice were trained to associate these artificial sensory cues with the rewarded direction and to indicate their decision by selecting the correct path leading to the rewarded port (Fig. 2d). Real-time location tracking and stimulation delivery were achieved using the EthoLoop tracking system (Nourizonoz et al., 2020), which ensured precise closed-loop operation with minimal latency (see Methods). All mice expressing ChR2 (n=6, 3 mice per maze, see Methods) successfully associated sensory cortex stimulation with the correct direction after visiting each intersection (Supplementary Video 1), while control mice (not expressing ChR2, n=5, 3 in the small maze and 2 in the large maze) performed at chance level (Fig. 2e, Supplementary Fig. 8).

**Figure 2:**
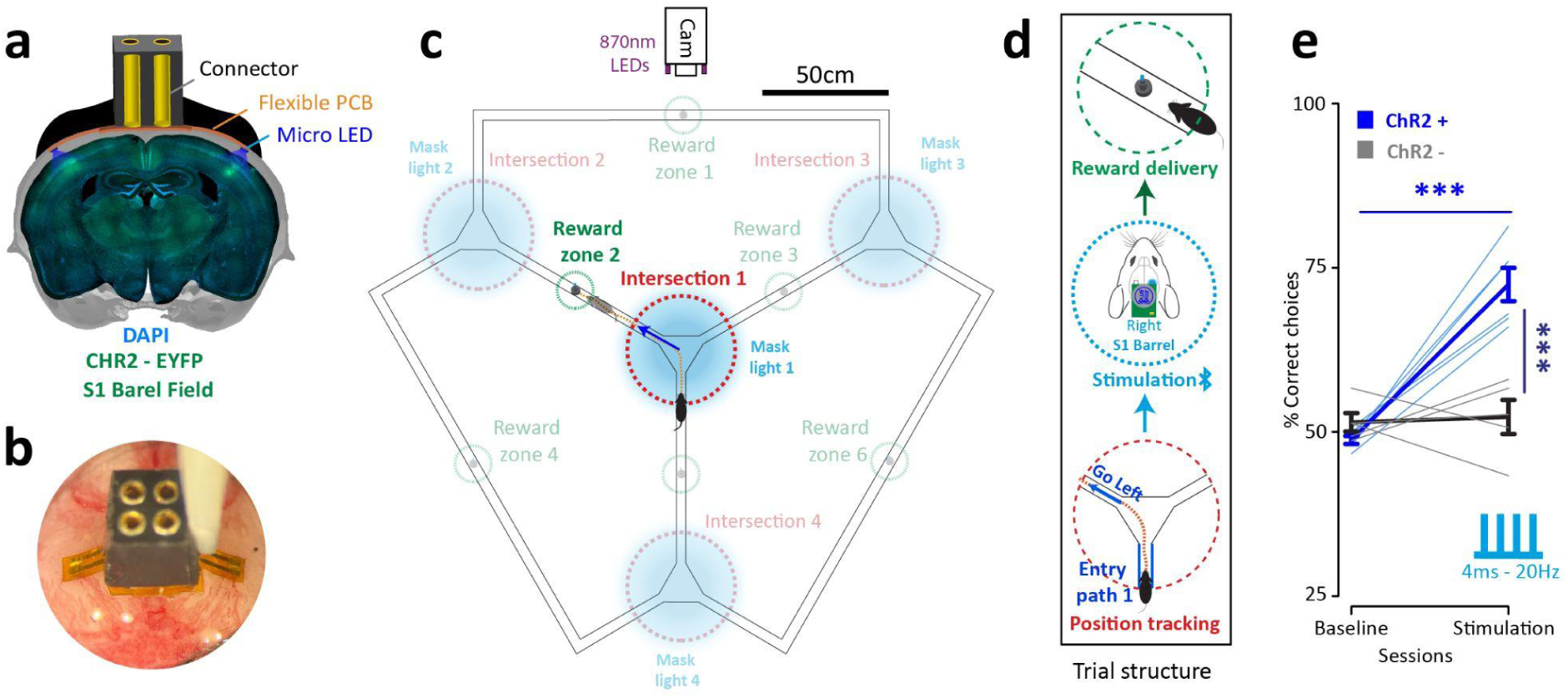
Closed-loop control of large-scale navigation using the BlueBerry. **a)** Coronal section of mouse brain with bilateral ChR2 expression on the barrel sensory field and the LED implant setup. **b)** Top view of the arrangement of the two-channel LED implant on the skull before covering with dental acrylic. **c & d)** Detailed configuration of the infinit Y-maze along with an example highlighting the trial structure. Mice arriving at the intersection zones received optogenetic stimulation based on a randomly chosen path (contralateral barrel sensory field) that leads to the arm containing a rewarded port. Reward is only delivered when mice choose the correct path and arrive in the corresponding reward zone. **e)** Performance of mice in first (no stimulation) versus the last session (4ms - 20Hz) to associate artificial sensory cues with the correct chosen path at every intersection in both small and large infinit Y-maze (*N* =6 ChR2 expressing mice, blue lines, 3 mice per maze; N=5 control mice not expressing ChR2, grey lines, 3 for small and 2 for large maze; two-way repeated measure ANOVA, main and interaction effect *P*< 0.01, pairwise two-sample *t*-test,Tukey post hoc test, \*\*\**P* < 0.001). Bars represent standard error of mean.

We also observed a brief, reproducible decrease in running speed following optogenetic stimulation of the barrel cortex in ChR2-expressing mice (Supplementary Fig. 9), an effect absent in control animals (Supplementary Fig. 9). To determine if this effect was related to the optogenetic cortical activation or to the decision-making process, we used BlueBerry to provide visible visual cues instead of optogenetic stimulation (Methods and Supplementary Fig. 10). Mice could reliably use visual cues to navigate but showed no speed change, suggesting the slowdown during optogenetic trials is not linked to decision-making processing (Supplementary Fig. 9). A possible explanation for such fast and reproducible motor effects caused by stimulation of the sensory barrel field could be the activation of direct descending pathways affecting the motor centers (Barsy et al. 2010).

Overall, these experiments show that BlueBerry can be seamlessly integrated into large-scale navigation paradigms, providing artificial somatosensory cues that shape decision making in freely moving mice.

### Transient shaping of social group dynamics

Complex social behaviors, such as dominance hierarchies, emerge from repeated interactions and are typically stable over time (Lindzey, Winston, and Manosevitz 1961; Wang et al. 2011; Fetcho et al. 2023). These hierarchies are expressed through asymmetric behaviors including chasing and aggression (Wang et al. 2011; So et al. 2015). Dopaminergic neurons in the ventral tegmental area (VTA-DA) are known to contribute to social reinforcement learning (Solié et al. 2022), raising the question of whether closed-loop stimulation of VTA-DA neurons can influence ongoing social interactions and transiently reshape group dynamics.

We used BlueBerry to reinforce specific social interactions which oppose the natural dominance hierarchy in groups of three male mice. DAT-iresCre mice (N = 9, housed in triads of siblings) were injected with a Cre-dependent ChR2 virus targeting VTA-DA neurons and implanted with fiber-coupled LED systems aimed at the VTA (Fig. 3a; Supplementary Fig. 5). We first confirmed the functionality of the implant and virus expression via closed-loop reinforced place preference experiments, where all subjects showed a significant preference for zones associated with optogenetic stimulation (Fig. 3b&c).

**Figure 3:**
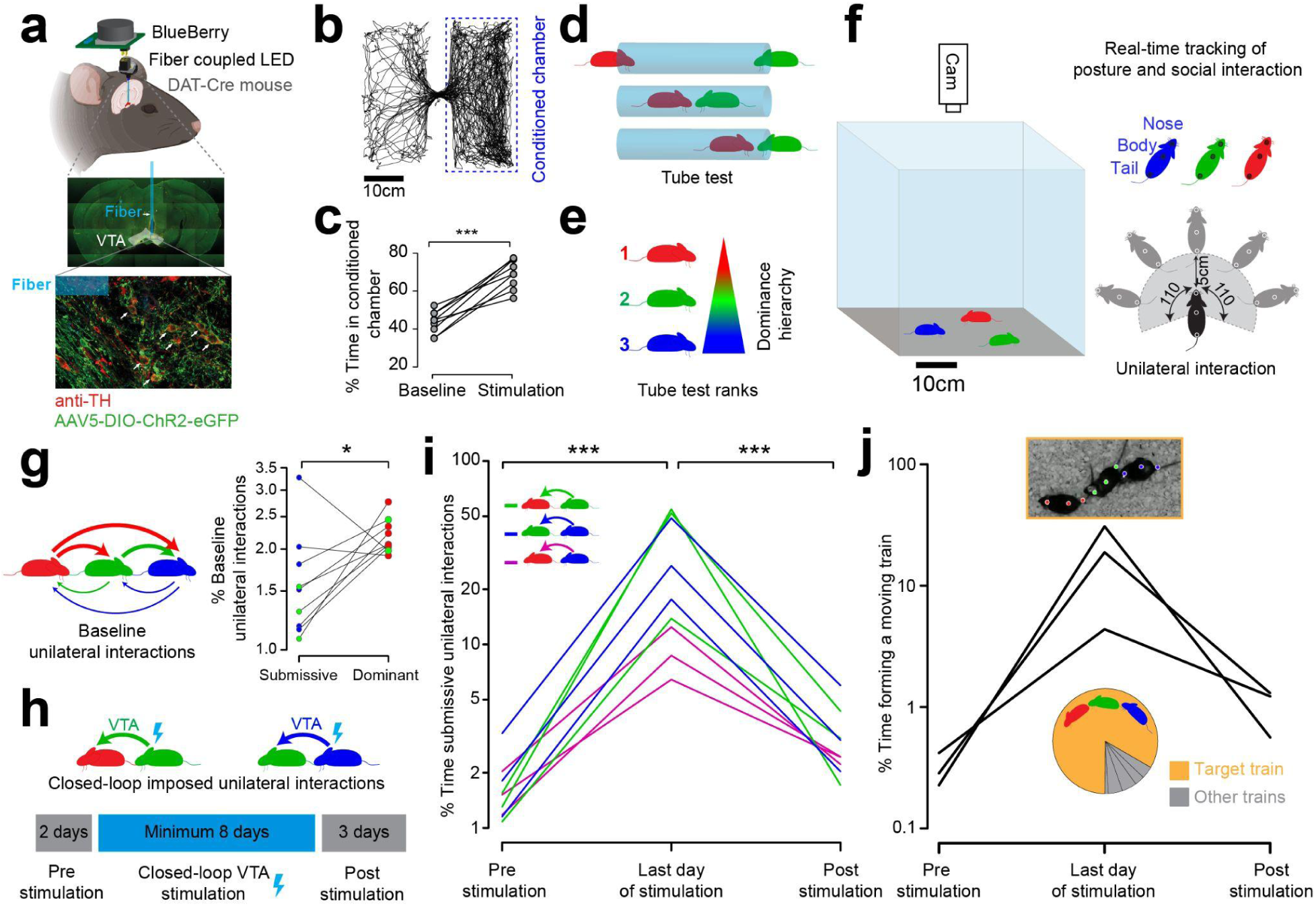
Closed-loop shaping of social behavior in groups of mice. **a)** Top: Fiber-coupled LED setup for freely moving mice to optogenetically stimulate DA neurons expressing ChR2 in the VTA. Bottom: Representative histology of the optical fiber trace and virus expression in the VTA region. **b)** Example trajectory of a mouse during place preference tasks (two chambers setting) where they received VTA optogenetic stimulation using the BlueBerry device upon entering a conditioned chamber. **c)** Performance of all mice in the real-time place preference task (N=9, pairwise student t-test, \*\*\**P*<0.001) **d)** Schematic of the tube test used to determine dominance hierarchy within each cage. **e)** Color coded hierarchy (ranks 1-3) for each cage based on tube test outcomes. **f)** Schema of real-time tracking and interaction detection. Simultaneous pose estimation of multiple freely moving mice is used to detect social interaction in real-time. Unilateral interaction is detected based on the angle (<110 degree) and distance (<5cm) of one mouse’s nose to another’s mouse tail. **g)** Right: Expected structure for baseline social interactions within a triad, with dominant interactions shown by thick arrows and submissive interactions by thin arrows. Left: Percentage of unilateral interactions initiated by submissive and dominant mice during baseline. The y-axis is scaled logarithmically (3 cages, 3 males per cage; pairwise t-test on log-transformed interaction durations, *P < 0.05). **h)** Behavior-triggered reinforcement strategy: rank 2 (green) and rank 3 (blue) mice received optogenetic VTA stimulation upon initiating unilateral interactions toward rank 1 (red) and rank 2 (green) mice, respectively. Bottom: Experimental timeline showing pre-stimulation, stimulation, and post-stimulation phases. **i)** Percentage of unilateral dyadic interactions initiated by the submissive member of each pair across pre-stimulation (first day), stimulation (last day) and post-stimulation (last day) sessions. The y-axis is scaled logarithmically (N=9 interaction, 3 interactions per cage, one-way repeated measure ANOVA, main effect \*\*\**P*<0.001, Bonferroni-corrected paired t-tests for post hoc test on log-transformed interaction durations, \*\*\**P* < 0.001). **j)** Train formation probability across pre-stimulation, stimulation and post-stimulation sessions. Each black line represents a cage. The y-axis is scaled logarithmically. The pie-chart represents the distribution of all possible trains (with the target train shown in orange) averaged across three cages during the optogenetic stimulation session. (Top) Image of multiple mice during an optogenetic stimulation session forming a moving train. The plotted dots on each mouse shows the location of their body parts (nose, body and tail) and the color indicates their natural hierarchy rank as shown in panel (e).

Next, we investigated the social rank of each animal by performing the tube test (Lindzey, Winston, and Manosevitz 1961) across pairs of cagemates (Fig. 3d-e, Methods). Subsequently, mice from the same cage were monitored in a large 0.5 x 0.5m open field arena (Supplementary Fig. 11) for 2 hours per day, during which the total duration of social unilateral interactions were quantified in real-time (Fig. 3f &g) by tracking their body parts with DeepLabCut -Live ((Kane et al. 2020), Methods).

To test whether the established hierarchy could be challenged by selectively reinforcing certain social behaviors, we optogenetically stimulated VTA-DA neurons in lower-ranked mice whenever they initiated unilateral interaction (chasing) toward a higher-ranked animal (Fig. 3h). Specifically, for each cage, we attached two BlueBerry devices to the implants of the mice in ranks 2 and 3. These devices delivered VTA-DA stimulation when rank 2 chased rank 1, and when rank 3 chased rank 2 (Fig. 3h; Supplementary Fig. 12). This reinforcement protocol was conducted over eight consecutive days (2 hours per day) in the open-field arena, followed by a three-day observation period without stimulation (post-stimulation sessions) to assess any lasting effects on the social interactions.

During the reinforcement phase, all chasing behavior initiated by the lower rank mice (including the ones not directly reinforced (rank 3 toward rank 1) significantly increased (Fig. 3i, N=9 mice, 3 interaction sets per cage, Supplementary Fig. 13). However, this effect was not sustained at post-stimulation sessions and did not impact the rank in the tube test followed after the reinforcement phase (Supplementary Fig. 13).

Strikingly, while our closed-loop protocol directly reinforced only defined dyadic interactions, it also induced emergent group behaviors. We frequently observed all three cage-mates forming “moving trains” in which multiple mice followed one another in close succession, typically led by the originally dominant animal (Fig. 3j; Supplementary Video 2). Thus, closed-loop reinforcement of specific unilateral interactions via VTA-DA stimulation transiently modulated group-level social dynamics without permanently altering rank.

These results establish, to our knowledge, the first demonstration of real-time closed-loop optogenetic manipulation in multiple freely interacting mice using a fully wireless system. They highlight the potential of BlueBerry for behavior-specific reinforcement and the study of social circuits in naturalistic multi-animal settings.

## Discussion

We present BlueBerry, a wireless, battery-powered optogenetic system that enables efficient, low-latency manipulation of neuronal activity in freely moving mice, including in large-scale environments and multi-animal social settings. BlueBerry’s modular architecture, with its standard connectors, detachable device, and interchangeable LED implants, provides substantial flexibility in targeting superficial cortical and deep subcortical structures without requiring modifications to the core electronics. This design simplifies integration into existing experimental pipelines and facilitates rapid adaptation to diverse paradigms.

Compared to other published wireless optogenetic systems (Kathe et al. 2022; Michoud et al. 2021; Li et al. 2022), BlueBerry combines several advantages: compact and lightweight form factor, openly available documentation, reliance on off-the-shelf components, and straightforward assembly without specialized facilities. These features lower the barrier to adoption, particularly for laboratories without dedicated engineering infrastructure. A key innovation is the BlueHub control unit (Supplementary Fig. 3), which unifies device management, stimulation programming, and interfacing with external behavioral frameworks via simple TTL inputs. Its graphical user interface eliminates the need for custom code to configure stimulation protocols or coordinate multiple devices, thereby reducing the time and effort needed to deploy closed-loop experiments.

One current limitation of BlueBerry is that light output is adjusted by changing the LED or by manually changing resistors on the circuit board. Incorporating programmable voltage or current regulators in future versions would allow automated power control from the same BLE interface, further expanding flexibility, for example, enabling dynamic switching between LEDs of different wavelengths and facilitating the use of multiple opsins with distinct spectral sensitivities.

By relying on onboard batteries rather than external power coils, BlueBerry removes the spatial constraints associated with inductive power-transfer systems and similar infrastructure (Parker et al. 2023). To this end, we successfully demonstrated BlueBerry’s ability to control navigation in large-scale mazes via closed-loop stimulation of the primary somatosensory cortex, across distances that exceed those commonly used in standard lab setups. In this task, mice learned to interpret artificial somatosensory cues as navigational instructions, offering a powerful paradigm to probe decision-making processes distributed over time and space (Harvey, Coen, and Tank 2012; Tseng et al. 2022).

Interestingly, optogenetic stimulation of primary somatosensory cortex reliably induced a rapid, transient decrease in locomotor speed in ChR2-expressing mice. This effect was not observed in control mice or when optogenetic stimulation was replaced with visual cues. Because the slowdown occurred with short latency relative to stimulation and preceded the time point at which trajectories diverged toward different choices, it is unlikely to reflect deliberation or decision-making per se (Supplementary Fig. 9–10). Instead, it suggests direct effects of cortical activation on the descending motor system, consistent with reports that barrel cortex as a classical sensory area, contributes to motor control and that descending projections participate in whisker retraction (Barsy et al. 2010; Karadimas et al. 2019). From an ethological perspective, such pathways may support rapid withdrawal or adjustment behaviors when whiskers encounter unexpected objects (Luis-Islas et al. 2022; Pancholi, Ryan, and Peron 2023).

We also demonstrated BlueBerry’s multi-device capabilities by implementing closed-loop optogenetic reinforcement in groups of freely interacting mice, targeting VTA-DA neurons during specific unilateral interactions that go against the established social hierarchy. This protocol selectively increased chasing initiated by lower-ranked animals and generated emergent “train-like” group formations, indicating that reinforcing selected dyadic interactions can transiently reshape group-level dynamics. However, the reinforced behavior did not persist in the absence of optogenetic stimulation and did not affect the initial dominance hierarchy (Supplementary Fig. 13). This could possibly be explained by the experimental structure, where mice are stimulated for only 2 hours per day in a specific arena, different from their home cage, where no stimulation is applied. Therefore, we assume that the reinforcement was strongly associated with the environment, and upon returning to their home cage, the dominant mouse resumed its dominant behavior over the other two cage mates. Future studies are needed to determine whether prolonged stimulation periods or alternative grouping strategies (mice from different cages) yield more durable modifications.

Taken together, our results highlight the effective integration of the BlueBerry system with various open source or commercial experimental frameworks (EthoLoop, DeepLabCut, and Ethovision) to investigate a broad range of behaviors, from sensory-guided navigation to social interactions. As neuroscience increasingly moves toward more naturalistic and ethologically grounded experimental designs, accessible wireless optogenetic tools such as BlueBerry will be crucial for linking neural activity to behavior in realistic settings. We anticipate that the open, modular design and wide applicability of BlueBerry will foster its broad adoption in systems and behavioral neuroscience.

## Acknowledgment

We thank all members of the Huber lab for their support and discussions. We thank Mario Prsa and Alastair Loutit for their comments on the manuscript and to Lorena Jourdain for technical assistance with VTA injections. We also thank Varta Microbattery for supplying the CP 1254 A4 coin batteries. This work was supported by the Swiss National Science Foundation (310030_184829 and 310030_215062).

## Contributions

A.N. and D.H. conceptualized the BlueBerry system. A.N designed and constructed the BlueBerry hardware, BlueHub hardware and brain implants. A.N. wrote all software for BlueHub and BlueBerry. A.N. ran all the maze navigation experiments. A.N. carried out the surgeries for micro-LED implantation for sensory cortex stimulation. A.N and B.G. ran all the social behavior experiments. A.N analysed all the data. D.H. and C.B. oversaw data analysis. B.G. carried out all the VTA surgeries for social experiments. B.G wrote the software for real-time social behavior classification. B.G. and M.G. performed all histological verifications. G.G. provided expertise and guidance for virus expression and surgeries. R.T. and S.P. provided guidance and contributed to PCB design and hardware assembly. C.B. and D.H. supervised the social behavior experiments. A.N. and D.H. wrote the manuscript.

## Material and methods

### Hardware

#### BlueBerry

The BlueBerry optogenetic device consists of a bluetooth low energy module (RN4871) with integrated antenna and an 8-bit AVR microcontroller (ATTINY-85-QFN). A surface mounted red LED (QT BRIGHTEK, QBLP595-R) is connected to the microcontroller for monitoring bluetooth connectivity status. To increase the output current, a 2-channel mosfet is used between the microcontroller and the 2x2 male (Mill-Max, 852-10-020) connector for LED implants. Micro-switch (C&K, PCM12SMTR) is placed between (in series) a small lithium rechargeable coin battery (VARTA, CP1254-A3) to power the device through a low dropout voltage regulator (MAX8888EZK33+TCT-ND). To be able to charge the battery and upload the base program on each BlueBerry device (RN4871 and ATTINY-85 initial configuration) a 10 pin micro-connector (Hirose Electric, BM23PF0.8-10DP-0.35V) is used as an interface between the BlueBerry device and the BlueHub control unit (Supplementary Fig. 3). All the resistors and capacitors used for the design are in 0402 small package format. All the components except the battery are placed on top and bottom of a custom made printed circuit board (PCB, 10.5 x10.5 x 0.6 mm, Supplementary Fig. 1 & 2) using soldering paste (TS991AX35T4) and fixed in a reflow oven. The board size and design is restricted not to cover any area below the antenna of the bluetooth module. (Supplementary Fig. 2) In the end, both poles of the battery are manually soldered to positive and negative pads on the PCB. Connection and transmission of data to the Blueberry device is achieved either through the BlueHub (Supplementary Fig. 3), or directly through an Linux-based computer (see software section below) using a USB bluetooth dongle (ASUS).

#### Brain implants

For bilateral stimulation of the sensory barrel field, two high-power micro LEDs (Light Avenue, LA-UB20FP1) are placed at the ends of a custom designed flexible PCB (3.5mm long on each side, Supplementary Fig. 5). A 2x2 female connector (Mill-Max) is placed via 4 through holes at the center of the PCB for connecting to the BlueBerry. For VTA stimulation, a high-power surface-mount LED (2.5 × 3.5 mm, TOP LED 2835 BL) was coupled to a 10 mm long optical fiber (400 µm diameter, 4.5 mm shielded, FP400URT, Thorlabs). LED–fiber coupling was achieved using miniature 3D-printed components, secured with UV-curable adhesive (Norland 61, Supplementary Fig. 5). To assess potential temperature changes produced by the superficial micro-LED implants, we imaged the LEDs using a thermal camera (120 × 90 pixels, UNI-T UTi712S) placed above a 36 °C background created with a laboratory hot plate (IKA, HS-7).

#### BlueHub

The BlueHub control unit (Supplementary Fig. 3 & 4) uses an Arduino UNO R4 (with Wi-Fi/BLE connectivity) at its core, providing a link to BlueBerry devices and generating triggers based on four digital inputs connected to external BNC female connectors (Adam Tech, BNC RA PCB 50). The Arduino module drives a 2.42-inch screen (ZJY OLED ,128x64 pixels) for the GUI, allowing users to assign specific BlueBerry devices to each BNC channel and customize the stimulation parameters. A rotary encoder switch (Bourn Inc, PEC11R-4220F-S0024) enables menu navigation parameter adjustments. Momentary tactile switches (C&K, PTS645VM83-2 LFS) next to each BNC allow manual testing of the assigned functions. The BlueHub also contains three BlueBerry connection stations with female micro connectors (BM23PF0.8-10DS). Each of these connection stations are connected to a battery charging module (LTC4054LES5-4.2); two stations serve as charging-only stations, and the third station functions for both charging and uploading the base program (initial configuration) onto the BlueBerry device (the programming station). The programming station is connected to the Arduino for SPI-based programming of the BlueBerry’s microcontroller and two non-inverting buffer units (74AHCT1G125GW,125) for RX/TX communication with the BlueBerry’s Bluetooth module. For uploading the base program into the BlueBerry unit, the Wi-Fi/BLE Arduino UNO R4 of the BlueHub is replaced by a standard Arduino UNO to accommodate the required programming protocols.

### Experimental setup

#### Infinite Y maze

To create an elevated infinite Y-maze, wood composites structures were cut in 7cm bins to form bridges between the intersections and reward zones (Supplementary Fig. 7). The intersections were made by cutting the same type of structure into equilateral triangles with side lengths of 28cm. Smaller equilateral triangles (7cm side length) were cut from each three corners of the intersection platforms to create a symmetric entry via each elongated bridge. For creating protection on the bridges and intersections, flexible black PVC at 4mm thickness were cut at 5 cm bins to create walls around both sides of bridges and the intersection platforms. The attachment of walls were done through custom made 3D printed clips. The entire maze was elevated 50cm using 12 pipes (Ostendorf Kunststoffe 110mm diameter) below every intersection platform, reward zones and turning points. Six holes (3 mm diameter) were drilled in the middle of every bridge to place reward ports. Liquid reward was delivered through solenoid valves (Lee Company, LHDA0531215H) connected to the output reward port using silicon tubes (TYGON, inside diameter 1.6mm). The digital output channels of a WiFi Arduino (Arduino MKR1010) were used to trigger an optocoupler-based circuit, which converted the Arduino’s low-voltage signals into 12V to activate the solenoid valves for reward delivery. Four high power blue LEDs (Roithners, ELJ-465-627) were placed above each intersection platform (attached to ceiling) to create a mask light during optogenetic stimulation. Mask LEDs were controlled through a high power PNP transistor (BC328-16) each connected to digital ports of another WiFi Arduino (Arduino MKR1010). In order to have a strong reflection of the mask light for mice navigating in the maze, the intersection platforms and 20 cm of each entry bridge were fully covered with mat white tape (Dc-fix adhesive). In the end to avoid mice exiting the maze all the parts of the maze were covered with transparent plexiglass (Plexiglas ROHM, 4mm thickness).

For the small infinite maze setup (Supplementary Fig. 7), black PVC material was used on top of a white wooden platform (1m x 1m) to create six corridors intersecting at 4 different locations. The fixation of the corridors were done through custom 3d printed inserts nailed to the wooden platform at intersection locations. The middle of each corridor was drilled to place reward ports and the same system as the big maze was used for controlling the reward delivery. In the last stage a 1m x 1m plexiglass (Plexiglas ROHM, 4mm thickness) is placed on top of the maze.

In both small and large maze setups EthoLoop tracking system (www.etholoop.org, (Nourizonoz et al. 2020)) was used to track the location of freely moving mice. A retroreflective (2 x 9mm) tape was attached to the BlueBerry carried by the mouse for passive location tracking. All wireless communication for the reward delivery system and mask light was done through a high speed wireless router (ASUS, AC 3100).

#### Multi-animal arena

The arena for shaping social dynamics experiments Supplementary Fig. 11) was built using five plexiglass (50 x 50cm, Plexiglas ROHM, 4mm thickness) placed at the surface and surrounding sides. The transparent arena was placed inside a custom made wooden box (0.8 x 0.8 x 0.9m) covered with soundproof foam on all six sides. The wooden box was illuminated at 25 Lux homogeneously via 100 white LEDs forming a ring around the box diffused by a sheet. A camera (Logitech c920 1080p HD) was fixed at the top having a 90 degree downward view to the center of the transparent arena.

### Software

#### BlueBerry base program

The BlueBerry requires two separate programs for initial and one-time configuration; one for its Bluetooth module (RN4871) and one for its microcontroller (ATTINY-85). To upload these programs, the Arduino UNO Rev4 on the BlueHub is temporarily replaced with a standard Arduino UNO, and the BlueBerry is attached to the BlueHub’s programming port (Supplementary Fig. 3). The upload process involves two steps. First, the Arduino UNO is placed into reset mode (via the dedicated programming mode switch on the BlueHub) so it can serve as a USB-to-UART bridge for the RN4871. Using ASCII command lines in an Ubuntu terminal, the RN4871 is programmed to adjust its connection parameters (e.g., latency and interval), indicate connection status, and communicate with the ATTINY-85 over the I²C protocol. Once the Bluetooth module is configured, the Arduino UNO is taken out of reset mode, and the Arduino IDE is used to program the ATTINY-85 via SPI. This microcontroller program decodes incoming 2-byte packets to determine which LED channel to activate and sets parameters for blinking frequency, pulse width, and the total number of pulses.

#### BlueHub programing and configuration

The BlueHub device is programmed via the Arduino IDE, which uploads both the control logic and the graphical user interface (GUI) onto the Arduino hardware at its core (Arduino UNO Rev4). The control logic is organized as a state machine that handles user input from the rotary encoder switch, push button presses, and the BNC input channels. An integrated OLED screen is driven over an I²C interface, enabling the GUI to display and manage all configuration options. Within this interface, the user assigns each BNC input channel to a specific BlueBerry device ID and defines the associated stimulation protocol (e.g., frequency, pulse width, and number of pulses). For example, a high-level digital signal on BNC 1 can be set to trigger a particular stimulation program on BlueBerry ID2. All settings are stored in the Arduino’s EEPROM, preserving the assigned device IDs and stimulation parameters even when the BlueHub is powered off. Upon restarting, the BlueHub automatically recalls these saved configurations, therefore maintaining consistent settings across multiple sessions. Communication with BlueBerry devices is achieved using the Arduino BLE library.

#### Closed-loop control of navigation in Y-mazes

Tracking the position of freely moving animals was done using the EthoLoop 2D tracking software (on Ubuntu 20.04 environment) applied on undistorted images. The distance of animals to the center of each intersection was calculated in real-time to detect if they have entered any intersections through any of the three entrances (20cm threshold for large maze and 10cm for small maze). Upon entering an intersection, a bluetooth message for optogenetic stimulation is sent to the blueberry through Gattlib library (https://github.com/labapart/gattlib). The stimulation direction (left or right) is chosen randomly through C++ random generator. Simultaneously a UDP packet is sent to the WiFi Arduino of the mask light controlling unit that activates high power blue lights at the same frequency of optogenetic stimulation. Upon making the correct decision (exiting the chosen arm), the corresponding reward port will be activated through a WiFi UDP packet sent to the reward controlling unit. The reward message contains the solenoid valve number and duration of opening. A predefined matrix indicates which port is associated to each arm of all intersections. At the end of each session a configuration file is stored containing all information about the optogenetic stimulation protocol. The two-dimensional data along with time (in milliseconds), crossed intersection number, stimulation (left or right), reward delivery (correct or incorrect) and trial number was stored in a text file.

#### Closed-loop control of social behavior

We implemented our closed-loop stimulation for social behavior in MATLAB, which interfaced with a Python script via a memory-mapped file. This configuration created a shared memory space, allowing efficient data transfer and synchronization between the two environments. Each captured 500×500px image was transmitted to this shared memory location, where DeepLabCut-Live performed real-time body-part tracking (nose, body and tail) for each animal, identified by a unique shaving pattern (Supplementary Fig. 12, Supplementary Video 2). The MATLAB script continuously accessed the coordinates from the shared memory to compute distances and angles between each animal’s nose and tail to detect unilateral social interactions. Once such an interaction was identified between desired pairs of mice and remained for 200ms, the MATLAB program triggered a PulsePal device to generate a high-level digital signal (5V), which was sustained until the detected unilateral interaction between the pair of mice ended. (https://github.com/sanworks/PulsePal/tree/master/MATLAB). This signal was connected to the BlueHub’s digital inputs (BNC1 and BNC2), which had been configured to activate and deactivate the assigned BlueBerry devices according to predefined stimulation protocols (Supplementary Fig. 12). For two-chamber real-time place preference experiments, EthoVision software (Noldus) was similarly integrated with the BlueHub, enabling automated optogenetic stimulation via the BlueBerry device based on the animal’s location in real time.

### Animals

#### Surgeries for sensory barrel field stimulation

Cre-dependent channelrhodopsin mice (Ai32, RCL-ChR2(H134R)/EYFP) were anesthetized using 2% mix of isoflurane and oxygen and placed under the stereotactic frame. They were administered with dexamethasone (Mephamesone, 20ul intramuscular), carprofen (Rimadyl, 2.5mg/1000gr, 50ul subcutaneous), buprenorphine (Temgesic, 0.1mg/1000g, 25ul intramuscular) and lidocaine (Rapidocain 10 mg/mL, 50 μL subcutaneous). At least 20 minutes later, a small incision was made in the scalp skin to gain access to the skull. Two separate Craniotomies were made bilaterally over the sensory barrel fields (1.5 mm posterior, 3.5mm lateral to bregma) and two viral injections (50nl each) of Cre virus (AAV5.CamKII0.4.Cre.SV40, dilution 1:50 with 2% fast green) were performed at depth of 300 um to target layer 2/3 cortex (this step was not performed for control animals). The flexible two channel implant was fixed on top of the skull with cyanoacrylate glue so that the micro LEDs will be placed inside the craniotomies. Afterward the implant was secured and covered with black dental cement to avoid leakage of the LED lights.

#### Surgeries for VTA stimulation

DAT-iresCre mice (Slc6a3tm1.1(cre)Bkmn/J) were used. Mice aged 6–8 weeks were anesthetized using a mixture of oxygen (1 L/min) and 3% isoflurane (Baxter). Following anesthesia, the mice were placed in a stereotactic frame. The surgical area was prepared by shaving and disinfecting the skin, followed by local anesthesia with 40–50 µl of 0.5% lidocaine. Bilateral craniotomies were performed at precise stereotactic coordinates over the VTA (lateral ±0.5 mm, posterior −3.2 mm from Bregma, and depth −4.25 ± 0.05 mm). A viral solution containing rAAV5-Ef1a-DIO-hChR2(H134R)-eYFP (Addgene, titer ≥ 4.2 × 10^12 vg per ml) was injected bilaterally into the VTA, with a volume of 500 nl per side, using a glass micropipette. The virus was allowed to express for 3–4 weeks post-injection. Subsequent to the viral expression period, the mice were implanted with fiber-coupled LED implants targeting the VTA. For precise targeting, the fibers were implanted unilaterally (right side) using the coordinates: ML +0.5 mm, AP −3.2 mm, DV −4.20 mm from Bregma. The fibers were then secured to the skull with dental acrylic mixed with carbon powder. Injections and implantation sites were confirmed post hoc.

#### Histology

Mice were anesthetized with 3% isoflurane and euthanized with i.p. pentobarbital injection (Escornarkon, 150mg/kg). They were then transcardially perfused using a peristaltic pump (ISM829, Cole-Parmer) with 0.01 M PBS, pH 7.4 for 2 min, and then 4% paraformaldehyde (PFA) in phosphate sodium buffer (PBS; pH 7.4) for 3 min. The brains were post-fixed at 4°C in PFA 4% for 48h, washed 3x15min in PBS and then fully transferred into PBS. The brains were cut in slices of 60µm thickness for sensory barrel field and 50µm thickness for VTA with a vibratome (VT1000S, Leica). For sensory barrel field, the slices were incubated with slight agitation at room temperature with a Hoechst solution (dilution 1:1000, Invitrogen, #33342) during 5min for fluorescent nuclear counterstaining and washed again with PBS. For VTA, brain slices were washed three times with 0.1 M PBS, then pre-incubated in PBS-BSA-TX buffer (10% BSA, 0.3% Triton X-100, 0.1% NaN₃) for 60 minutes at room temperature in the dark. Slices were then incubated overnight at 4°C with primary antibodies diluted in PBS-BSA-TX (3% BSA, 0.3% Triton X-100, 0.1% NaN₃). The next day, they were washed three times with PBS and incubated for 60 minutes at room temperature with secondary antibodies in PBS-Tween (0.25% Tween 20). The primary antibody used was rabbit polyclonal anti-tyrosine hydroxylase (1:500, Abcam, ab6211), and the secondary antibody was donkey anti-rabbit 555 (Alexa Fluor, 1:500). The sections were mounted on Superfrost microscope glass slides (Epredia) using mounting medium (Fluoromount, Southern Biotech), coverslipped with 24x50 mm coverslips (Menzel-Gläser) and kept in dark at 4°C until further imaging. All photomicrographs were taken using a Zeiss Axio Scan.Z1 (Zoom x10) for sensory barrel field and Zeiss LSM700/LSM800 confocal microscopes for VTA.

#### Ethics

All mouse experiments and surgeries were conducted in Geneva, Switzerland, and received approval from both the local ethics committee and the Geneva canton authorities. The mice were housed in the animal facility, maintained at a temperature of 21°C and a humidity level of 50%. In each cage, no more than five mice were accommodated.

### Experimental procedure

#### Closed-loop control of navigation in Y mazes

Three weeks after the surgeries, animals’ access to water was removed and limited to 1ml per day. For both small and large mazes, animals performed two baseline sessions with no optogenetic stimulations. Each session contained 200 trials and only one session per mouse was performed each day. A trial consisted of animals entering an intersection, receiving a stimulation (left or right sensory barrel field), choosing an exit path and finally arriving at the water port. When mice selected the correct exit at each intersection, a loud click noise was produced, using a solenoid striking a plastic surface located outside the maze. This serves as an early indicator of their decision’s outcome and facilitates the learning process. In the initial sessions (the first 8 sessions), this auditory cue allowed mice to reconsider their choices (if they entered an incorrect arm, they could return to the intersection and make a different choice, triggering the click noise). After the 8th session, however, this sensory feedback was removed, preventing the animals from altering their decisions.

In the small maze animals underwent 7 sessions with optogenetic stimulations delivered at 20Hz. The pulse number and pulse width varied across the seven days starting from weak stimulation 4 pulses of 1 ms and going up to 6 pulses of 5ms width and finally down to 4 pulses with 4 ms width (Supplementary Fig. 8). To characterize the effect of optogenetic stimulation on speed in more detail, animals in big maze underwent a total of 20 stimulation sessions with early sessions (first 7 stimulation sessions) including weak optogenetic stimulations (1 pulse with width of 1 and 2ms, Supplementary Fig. 8 & 9). After the 7th day the stimulation became stronger in terms of pulse number and pulse width and followed the exact same trajectory as the stimulation protocol used for the small maze (4 pulses of 4ms in the last day). All mice from the large maze experiment underwent 6 additional sessions where optogenetic stimulation was replaced with visual cues provided via two channel LED headstage (fabricated via 3D printer, 4 pulses of 4ms delivered at 20Hz, Supplementary Fig. 10). In both small and large mazes, animals were supplemented with additional water if they did not receive 1ml during the training session.

#### Shaping social dynamics

Three experimental cages, each housing three DAT-iresCre mice (Slc6a3tm1.1(cre)Bkmn/J) that had been cohabiting since birth were used. Three weeks post fiber-coupled LED implantation surgery, these mice were subjected to an optogenetic real-time place preference (rtPP) experiment, to validate the effectiveness of virus injection and the LED-fiber setup for optogenetic stimulation of dopaminergic neurons in the ventral tegmental area (VTA-DA). The rtPP experiment was conducted in a two-chamber setting. It began with a fixed 5-minute baseline session, during which the mice’s chamber preference was determined. This was followed by a stimulation session lasting 15 minutes. The less preferred chamber, as identified during the baseline, was designated as the conditioned chamber for the stimulation sessions. Within this chamber, VTA-DA stimulation was provided as a continuous 20 Hz train of 5 ms pulses.

After the rtPP experiment, the dominance hierarchy within each cage was determined using a tube test. This involved releasing pairs of mice at opposite ends of a transparent plexiglass cylinder (30 cm long and 3 cm in diameter), with the trial finishing when one mouse exited the tube. The mouse remaining in the tube is considered the dominant of the pair. This test was repeated three times daily for each pair, resulting in a total of nine trials per cage per day, over three stable consecutive days. The mouse winning all tube tests over two other mice is considered as dominant of the cage (rank 1), mouse winning tube tests against only one animal is considered as the middle in hierarchy (rank 2) and finally mouse losing all tube tests is considered as the submissive of the cage (rank 3). For identification during tracking, each mouse was shaved according to its hierarchical status; the submissive on the lower body, the middle near the neck, and the dominant remained unshaven (Supplementary Fig. 11 & 12). Initially, two baseline sessions (one session per day, 120 min) for each cage were performed where images of freely moving mice interacting with each other in the social arena were recorded. These images were used to train DeepLabCut model for real-time tracking purposes. Once the baseline was completed, each cage underwent eight sessions (8 days, 120min per day) where VTA-DA optogenetic stimulation was only delivered (continuous train of 5ms pulses at 20Hz) to the Rank 2 and Rank 3 mice only if engaging in a unilateral interaction toward the mouse one level above them in the dominance hierarchy (Rank 3 -> Rank 2 and Rank 2 -> Rank 1). Unilateral interaction was identified by monitoring the real-time positioning of the animal’s body parts, including the nose and its angular relation to another mouse’s tail. This type of interaction is confirmed when the nose-to-tail angle falls below 110 degrees, within a proximity of 5cm, and lasts at least 200 milliseconds. To evaluate the post effects of optogenetic stimulation, a tube test and three additional baseline sessions were conducted post-stimulation, enabling the observation of both changes in hierarchical status and social behavioral changes in the mice.

### Post hoc data analysis

#### Navigation in Y-maze

All data analysis and statistics were performed in the R (Rstudio) environment. Data processing includes smoothening of the positional data (10ms window) and creating a separate log file containing all delivered stimulation and the related performance of the mouse for any given stimulation at every intersection. The first 20 trials were excluded for all animals as they exhibit exploratory behavior in the maze and performance was measured across the next 150 trials. Comparison between control and channelrhodopsin positive mice were made based on their performance on the second baseline session and the last stimulated session (7th session in small maze and 24th session in large maze). To extract the speed from position data, a 40 ms sliding window was applied on the two dimensional position data. Speed comparison across different trials (left, right and blank) were made by aligning all the speed profiles to the stimulation point (time zero). For normalization, all speed values were divided to the mean of speed during optogenetic stimulation.

#### Social behavior

All position data (nose, body and tail) were initially interpolated at 40 ms intervals in the MATLAB environment and further smoothed in the R programming environment. Afterward, the interactions between pairs of mice (unilateral) were assessed again on the processed data. All statistical analysis and plotting were performed in R.

