## Supplementary Figures 1-13 for "BlueBerry: Closed-loop wireless optogenetic manipulation in freely moving animals"

### Supplementary figures and material

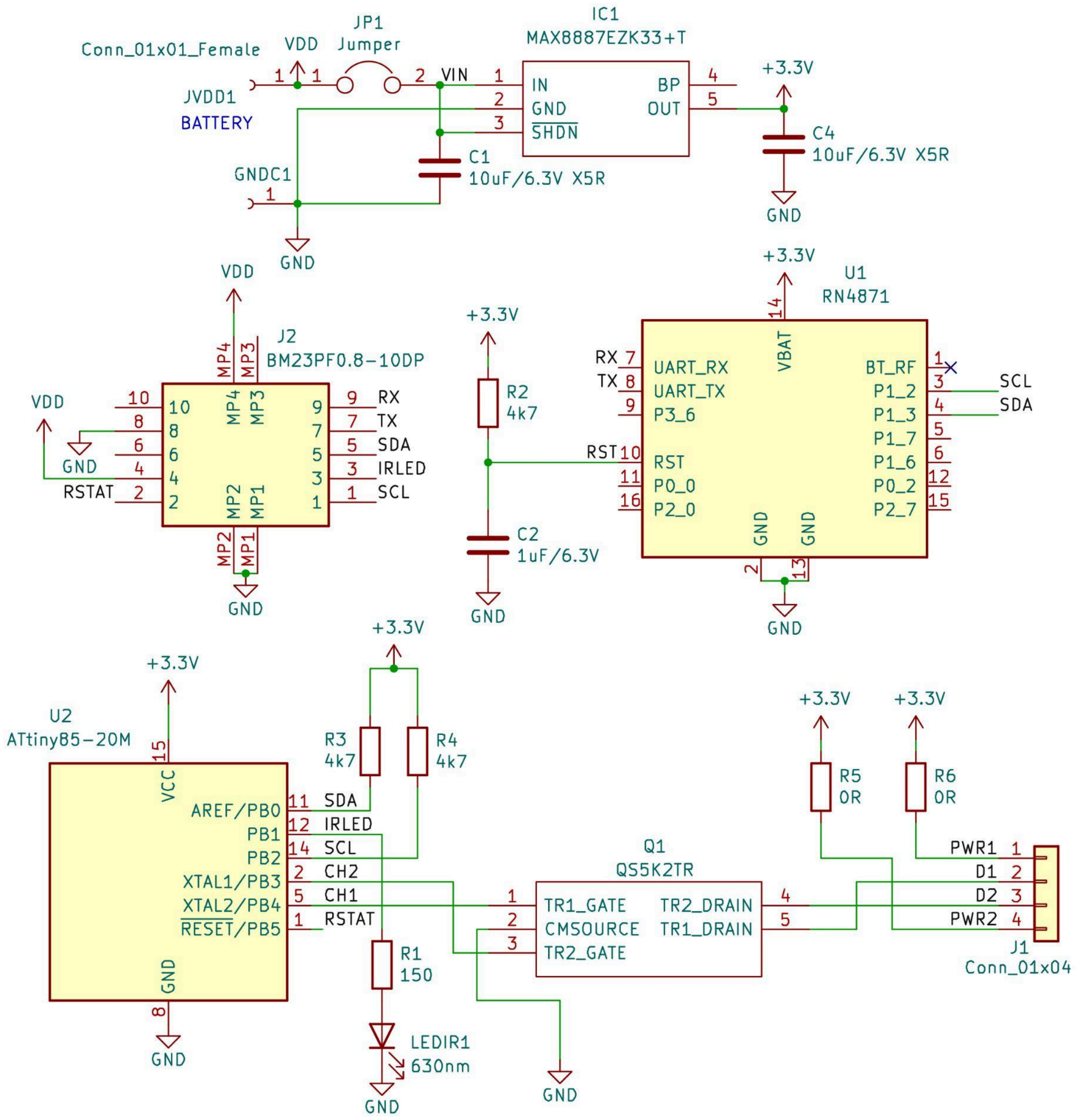


**Supplementary Figure 1: BlueBerry electronic schematic.** Detailed schematic of the BlueBerry device with all electrical components.

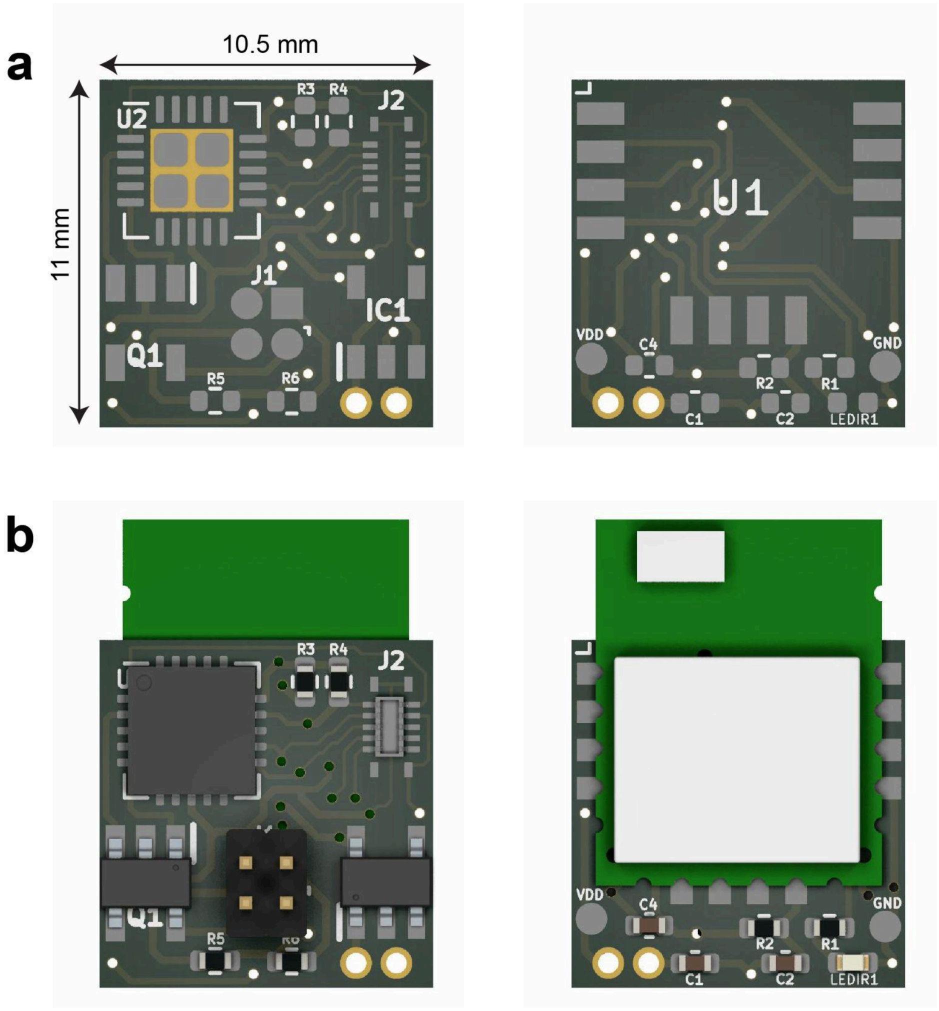


**Supplementary Figure 2: BlueBerry PCB.** Back and front side of the PCB (from left to right) layout of the BlueBerry device (top) and the mounted version on both sides without the micro switch and coin battery (bottom)

**Supplementary Table 1. Electrical components for assembling the BlueBerry device**

| Component Label | Description | Name/Value |
|---|---|---|
| J1 | 2 Channel LED Output | MillMax 1.27mm connector (M) |
| J2 | Micro Connector | BM23PF0.8- 10DP |
| Through Hole | Micro Switch | C&K Slide SPDT |
| U1 | BLE Module | RN4871 |
| U2 | Microcontroller | Attiny 85 - QFN |
| Q1 | 2 Channel Mosfet | QS5K2TR |
| IC1 | Voltage regulator | MAX8887EZK33+T |
| LED | Micro LED 0402 | Red LED |
| R1 | Resistor 0402 | 150 Ω |
| R2 & R3 & R4 | Resistor 0402 | 4k7 Ω |
| R5 & R6 | Resistor 0402 | **Adjust for your LED and power.** (0 Ω for high power blue LED) |
| C1 & C4 | Capacitor 0402 | 10uF /6.3V |
| C2 | Capacitor 0402 | 1uF /6.3V |

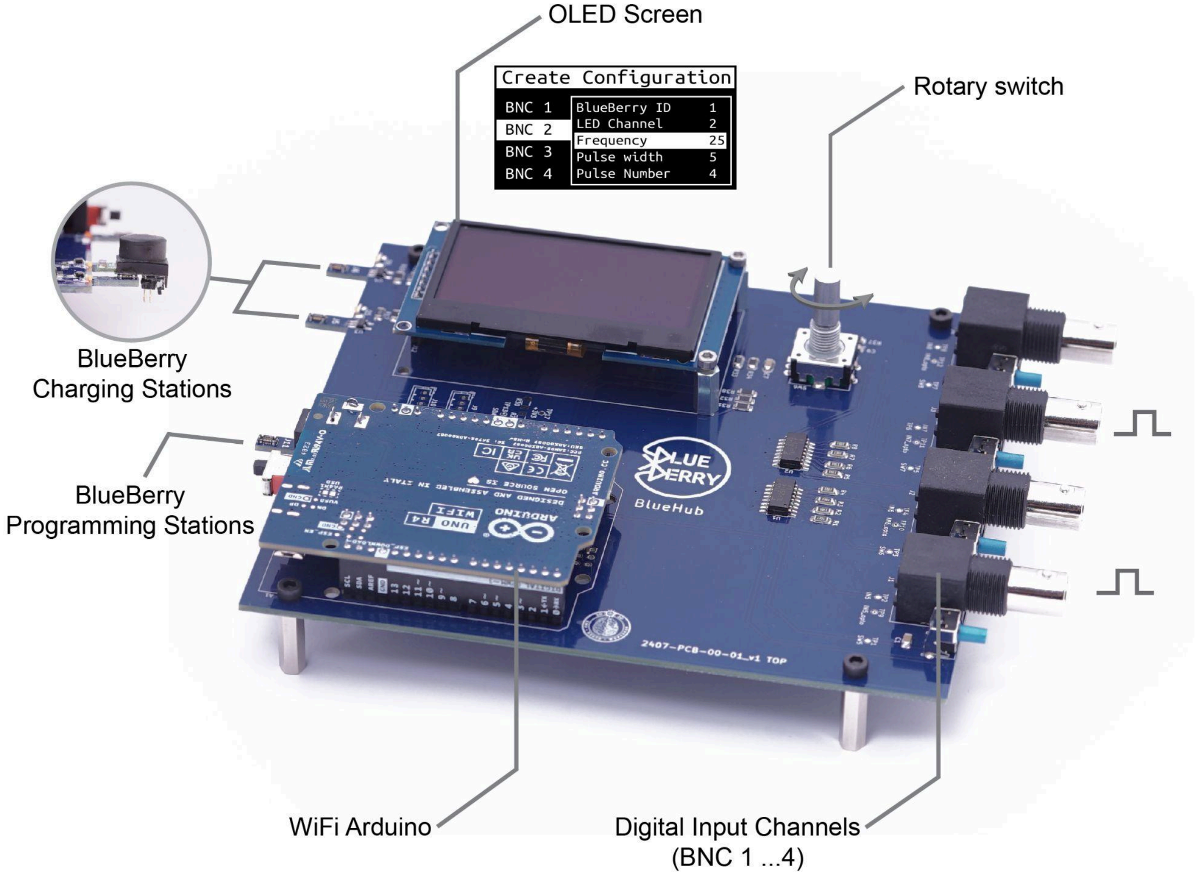


**Supplementary Figure 3: BlueHub control device.** The BlueHub control device is built around a Wi-Fi/BLE Arduino UNO Rev4, which communicates with BlueBerry devices in real-time. Through a dedicated GUI displayed on the integrated OLED screen, users can assign specific BlueBerry devices and stimulation protocols to each of four TTL input channels, enabling smooth integration with various behavioral frameworks for closed-loop interaction. Menu navigation and parameter selection are carried out via a rotary encoder. Two built-in charging stations with micro connectors provide convenient recharging of BlueBerry batteries after experiments. A third station allows base program uploads (initial Bluetooth and microcontroller configuration) to new BlueBerry devices by temporarily replacing the Arduino UNO Rev4 with a standard Arduino UNO.

a

b

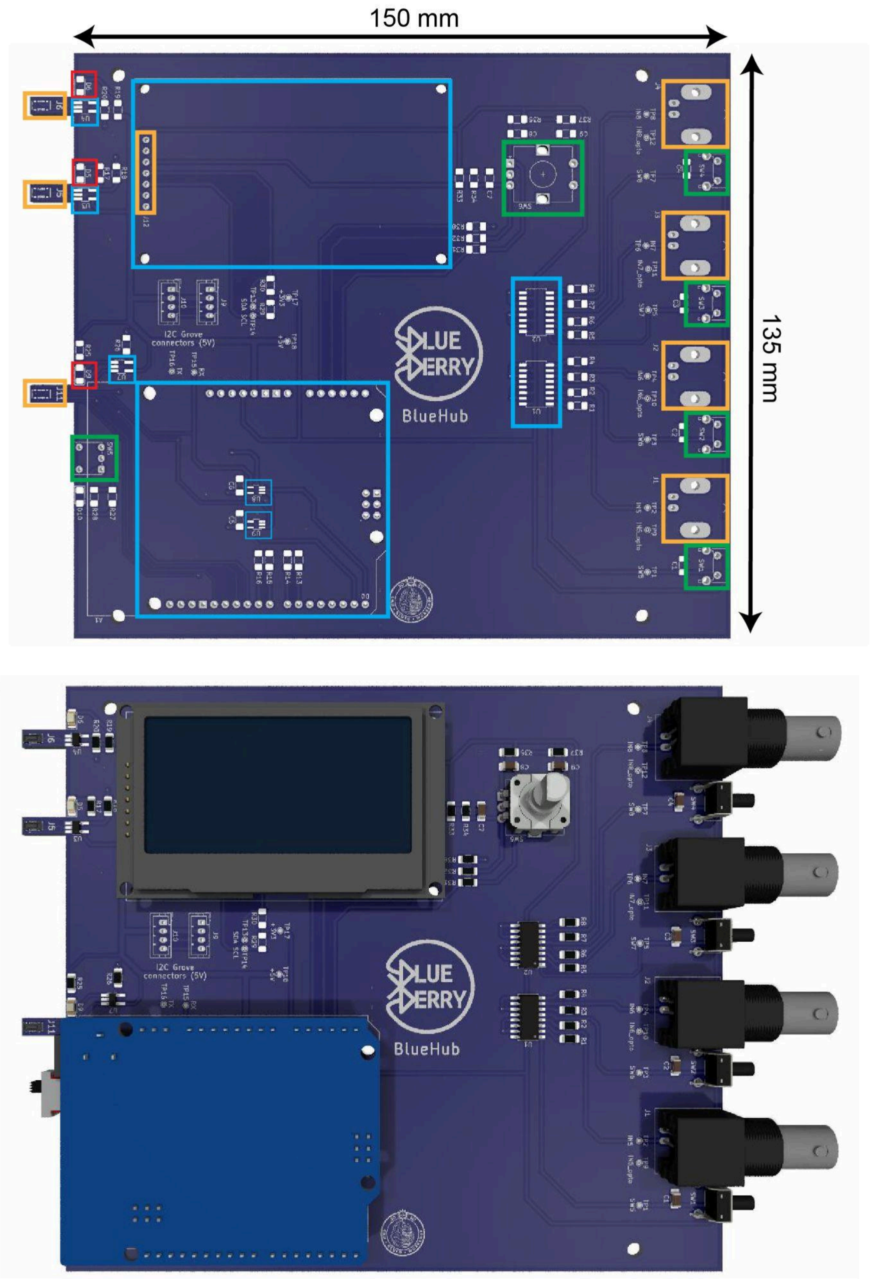


**Supplementary Figure 4: BlueHub PCB.** The PCB layout of the BlueHub device (top) and the mounted version of the PCB (bottom). The colors on the top PCB layout correspond to elements' type indicated in Table 2. Orange for through hole connectors, Green for mechanical switches, blue for integrated circuits and red for LEDs.

**Supplementary Table 2. Electrical components for assembling the BlueHub device**

| Component Label | Description | Name/Value |
|---|---|---|
| J1 to J4 | Female Right Angle BNC | Adam Tech |
| J5,J6,J11 | Female Micro-connector | BM23PF0.8-10DS-0.35V(895) |
| J12 | Screen connector | Arduino Stackable Header kit |
| SW1 to SW4 | Right Angle Switch | Tactile SPST-NO 0.05A 12V |
| SW5 | Right Angle Switch | Slide SPDT 3A 120V |
| SW6 | Vertical Switch | Rotary Encoder |
| U1, U2 | Optoisolator 3.75k | TLP293-4 |
| U3,U4,U7 | LI-ION Battery Charger | LTC4054LES5-4.2 |
| U8, U9 | Buffer, Non-Inverting | IC BUF 5.5V 5TSSOP |
| Screen at J12 | OLED screen | 2.42 inch ZJY screen, 128x64 |
| A1- Arduino | Wifi & BLE Arduino | Arduino UNO R4 |
| D1 to D4 (Back side of PCB) | Right Angle SMD LED | Blue LED, 3x2x1mm |
| D10 | Right Angle SMD LED | Orange LED, 3x2x1mm |
| D5, D6, D9 | Right Angle SMD LED | Red LED, 3x2x1mm |
| R1 to R8 | Resistor 1206 | 1.3k Ω |
| R13 to R16<br>R31, R32, R34, R35, R37, R38<br>R20, R18, R26, R28 | Resistor 1206 | 10k Ω |
| R9 to R12 (Back side of PCB) | Resistor 1206 | 5.9 Ω |
| R29 & R30 | Resistor 1206 | 2.2k Ω |
| R33 | Resistor 1206 | 4.7k Ω |
| R19, R17, R25 | Resistor 1206 | 150 Ω |
| R27 | Resistor 1206 | 59 Ω |
| C1 to C6 | Capacitor 1206 | 100nf |
| C7 to C9 | Capacitor 1206 | 10nf |

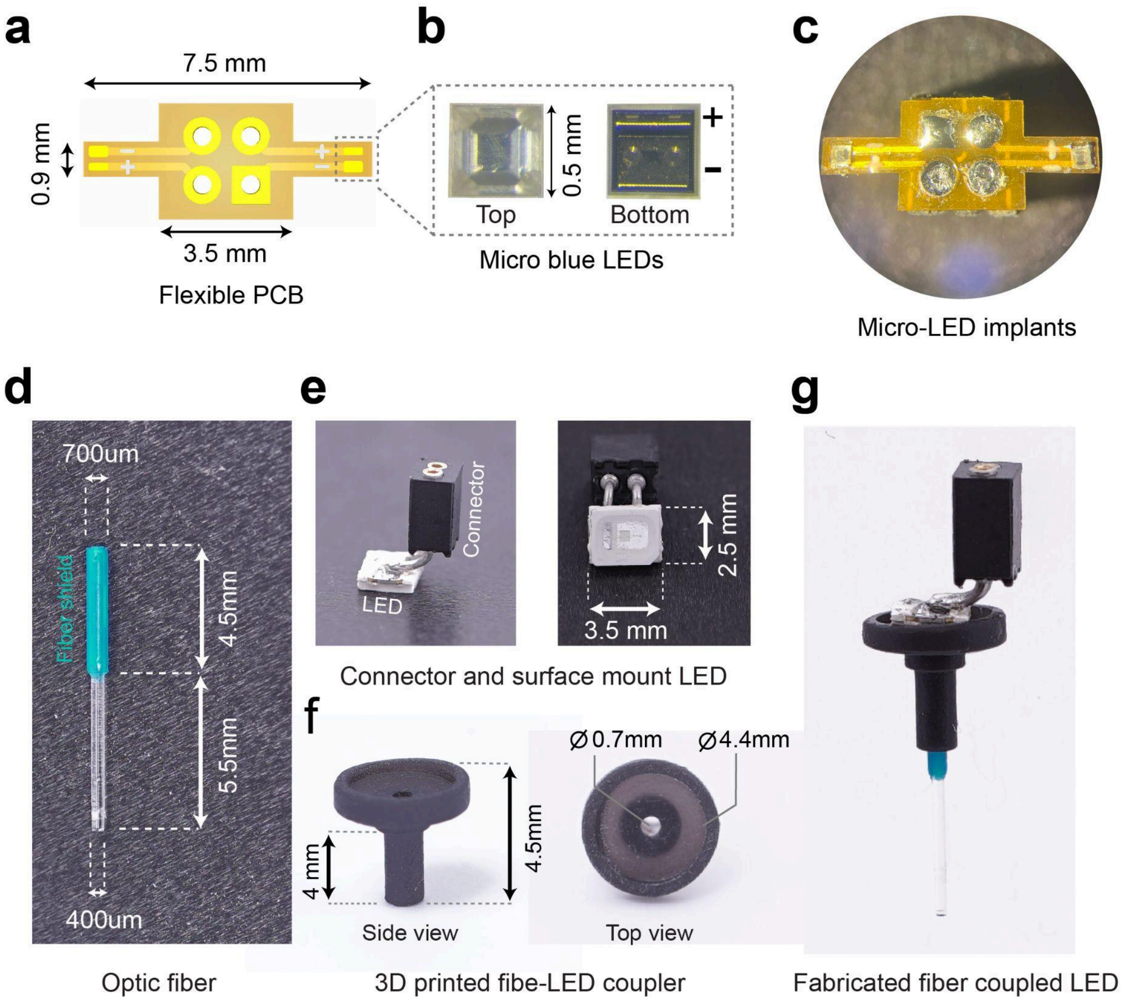


**Supplementary Figure 5. BlueBerry brain implants. a)** Flexible PCB layout of the multichannel micro LED implant for bilateral stimulation of the barrel sensory field. **b)** Detailed picture of top and bottom view of micro LEDs. **c)** Top picture of the micro-LED implants under microscope. **d)** Optic fiber preparation for targeting the VTA through coupling it with a high power LED. **e)** The surface mount high power LED setup soldered to a female 2x1 connector. **f)** Picture of the 3D printed element that couples the fiber shown in (a) with the LED-connector setup shown in (e). **g)** Side view of the final single-channel fiber-coupled LED implant for optogenetic stimulation of deep brain structures in freely moving mice.

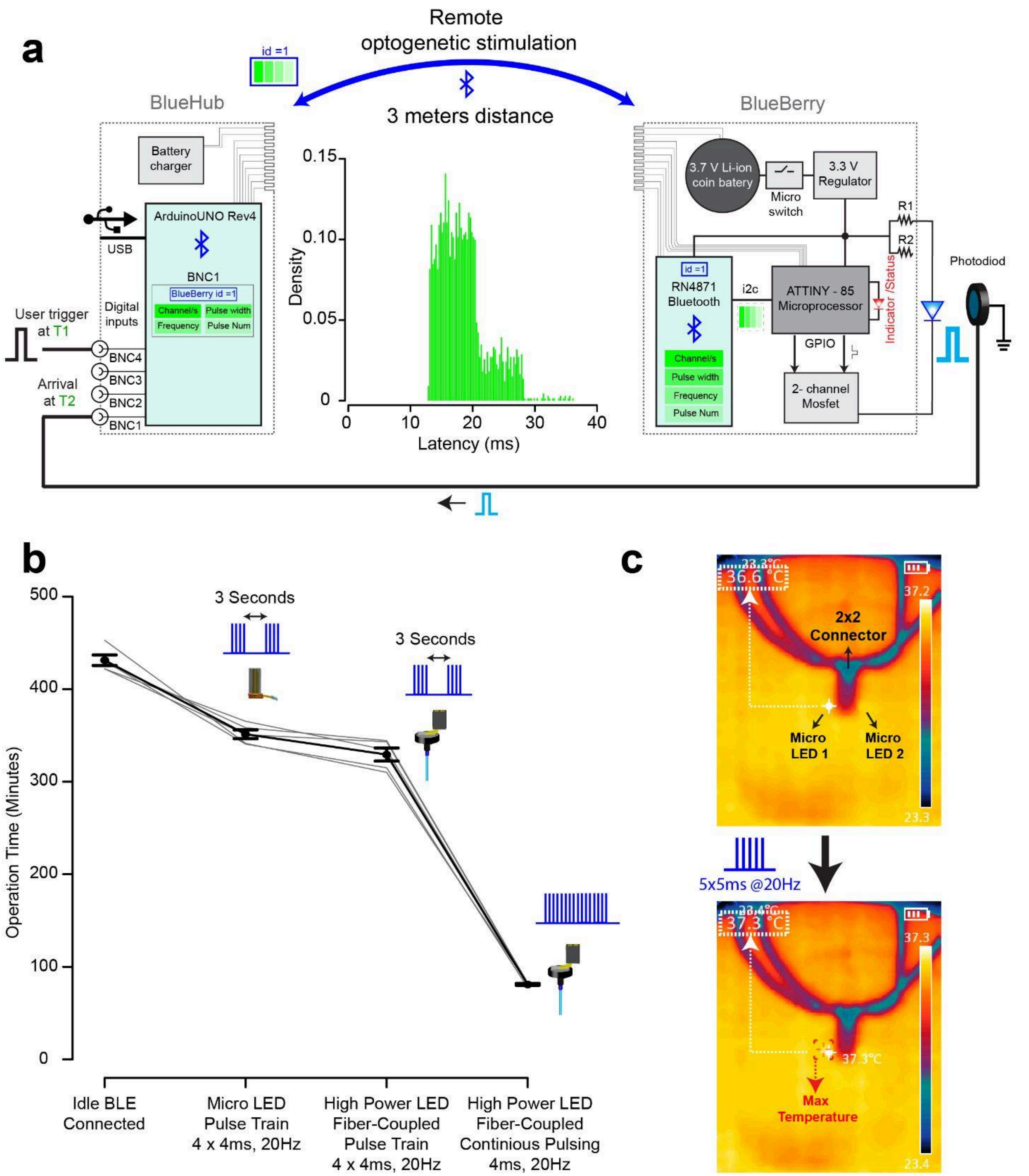


**Supplementary Figure 6. BlueBerry performance characteristics** **a)** Measurement of BlueBerry communication latency. To quantify the time delay between stimulus command and LED activation, one BlueHub input channel was used to trigger a BlueBerry device located 3 meters away, recording the trigger time as T1. Upon receiving the command, the BlueBerry device illuminated an LED positioned in front of a photodiode detector, which relayed a digital pulse back to another BlueHub input channel, logged as T2. The communication latency was defined as T2 – T1. This procedure was repeated 1,000 times, and the resulting latency distribution is shown as a histogram. **b)** Operation time of the BlueBerry device in four conditions. Idle with active wireless connection only; 3 s–spaced pulse trains via micro-LED implants for cortical stimulation; 3 s–spaced pulse trains via a fiber-coupled LED for deep/subcortical stimulation; and continuous 20 Hz stimulation via the fiber-coupled LED (N=5 BlueBerry devices). Bars represent mean ± s.e.m. **c)** Temperature measurements of micro-LEDs used for superficial/cortical stimulation. Top: thermal images of the bilateral micro-LED implant mounted on a hot plate set to ~36.5 °C (to simulate brain temperature), before and immediately after delivery of 5 pulses (5 ms, 20 Hz) using the same protocol as in the maze experiments. Temperature at the LED site (white cross) is indicated in the top-left corner of each image. The measured temperature increase (0.6 °C) is below 1 °C and thus complies with ISO 14708-1:2014(E) safety guidelines.

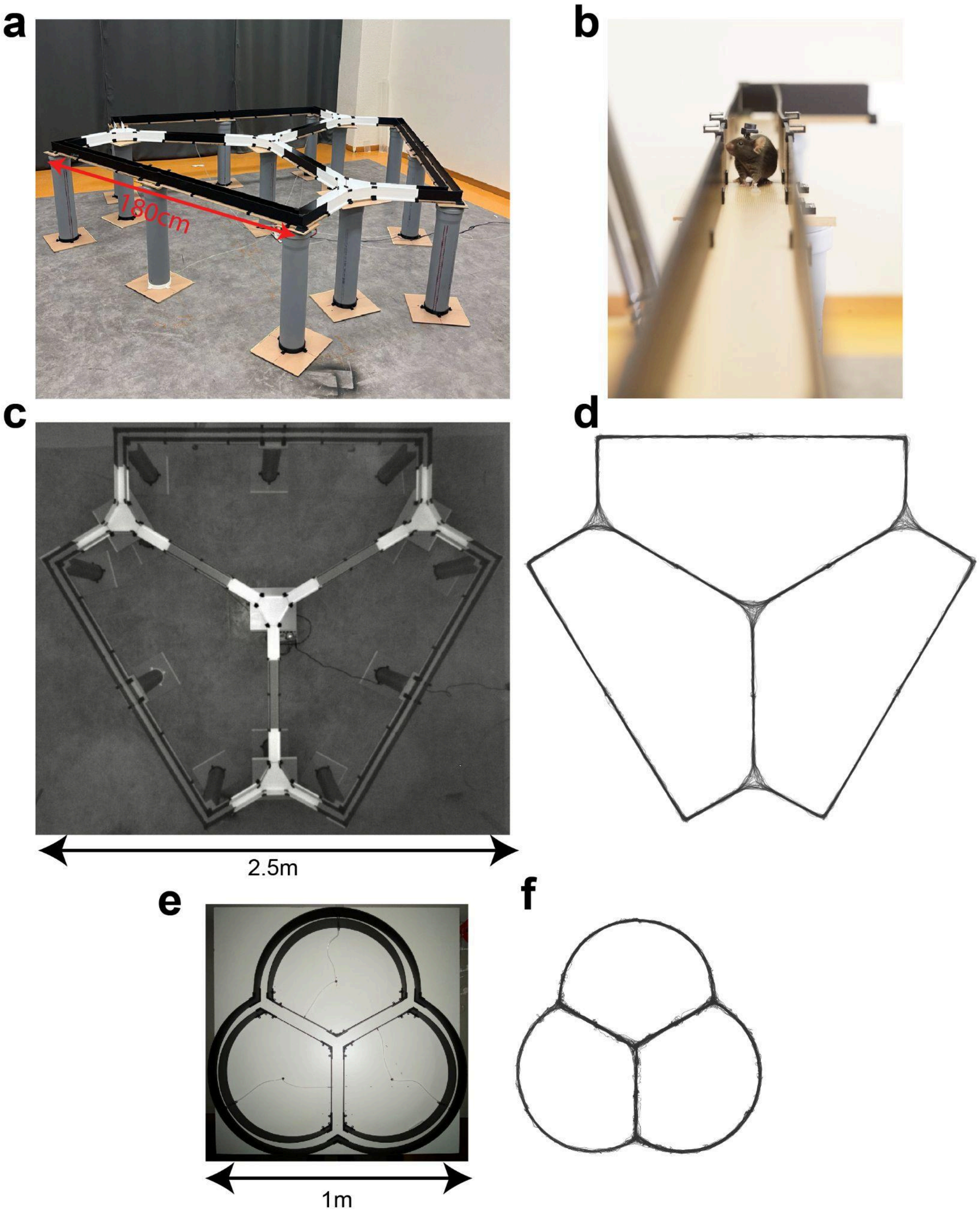


**Supplementary Figure 7: Experimental setup for infinit Y maze navigation task. a)** Picture of the elevated large infinit Y-maze. Intersection points are covered with mat white tape to reflect the blue mask light during optogenetic stimulation. **b)** Picture of a mouse with a BlueBerry device in one of the arms of elevated large infinit Y-maze. **c)** Tracking camera view of the large infinite Y-maze. The images of the tracking camera are undistorted in real-time to achieve correct and calibrated tracking. **d)** 20 minute tracking data of a mouse navigating inside the large infinit Y maze. **e)** Picture of the small infinit Y-maze. **f)** 20 minutes trajectory of a mouse navigating inside the small infinit Y maze.

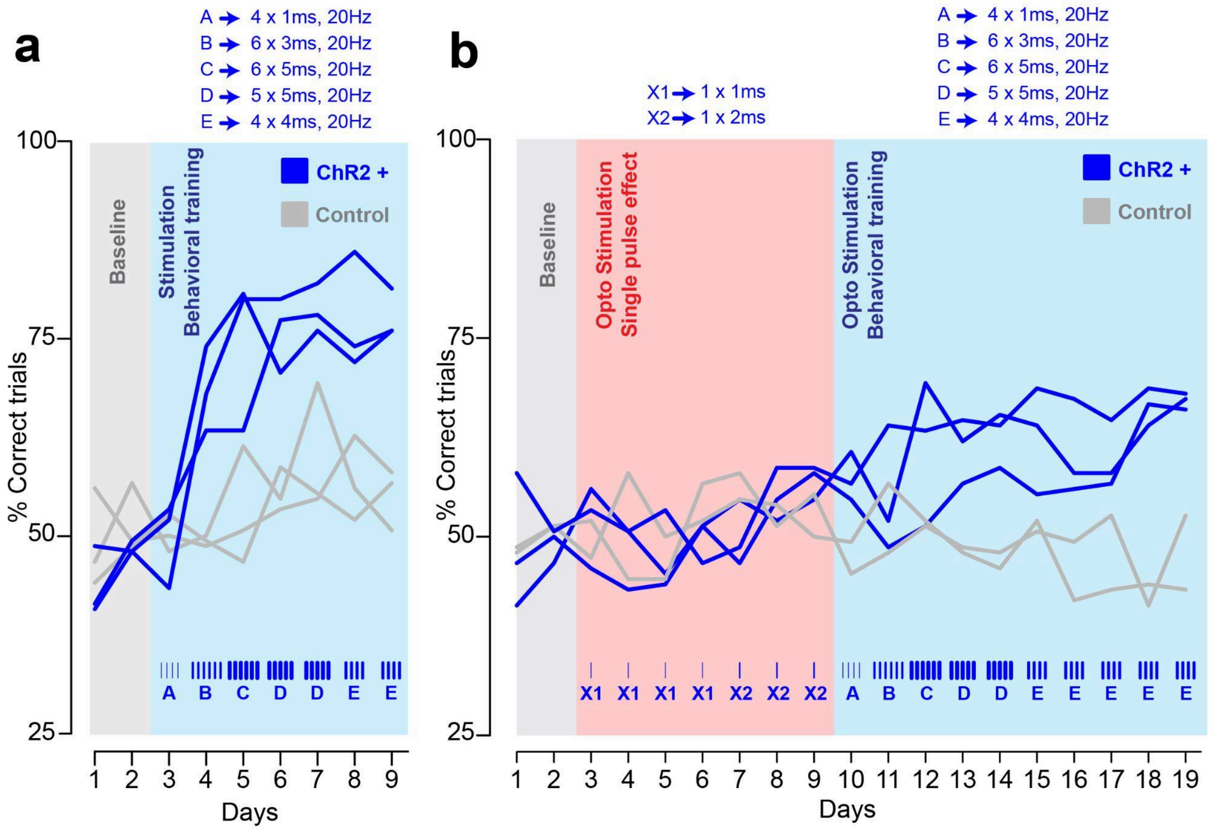


**Supplementary Figure 8. Training procedure and animal performance in infinite Y-maze navigation a)** Performance of 6 mice in the small maze (3 ChR2+ and 3 ChR-). Two baseline sessions (grey, no stimulation) were followed by 7 sessions with optogenetic stimulation at the maze intersections. Top: overview of the five stimulation patterns (A–E). Blue bars indicate the stimulation pattern per day (bar thickness = pulse width; number of bars = number of pulses). Letters (A–E) below the x-axis indicate the pattern used and correspond to the schematic above. **b)** Performance of 5 mice in the large maze (3 ChR2+ and 2 ChR-). 2 baseline sessions were followed by 7 sessions where only 1 pulse of optogenetic stimulation was delivered upon arriving at the intersections. This phase was mainly performed to characterize motor responses to optogenetic stimulation of primary sensory cortex in freely moving mice. Afterward, mice underwent 9 sessions of optogenetic stimulation with stimulation protocols identical to the ones provided in the small maze but the last day (day 7) was repeated twice. Letters below the patterns correspond to those in (a).

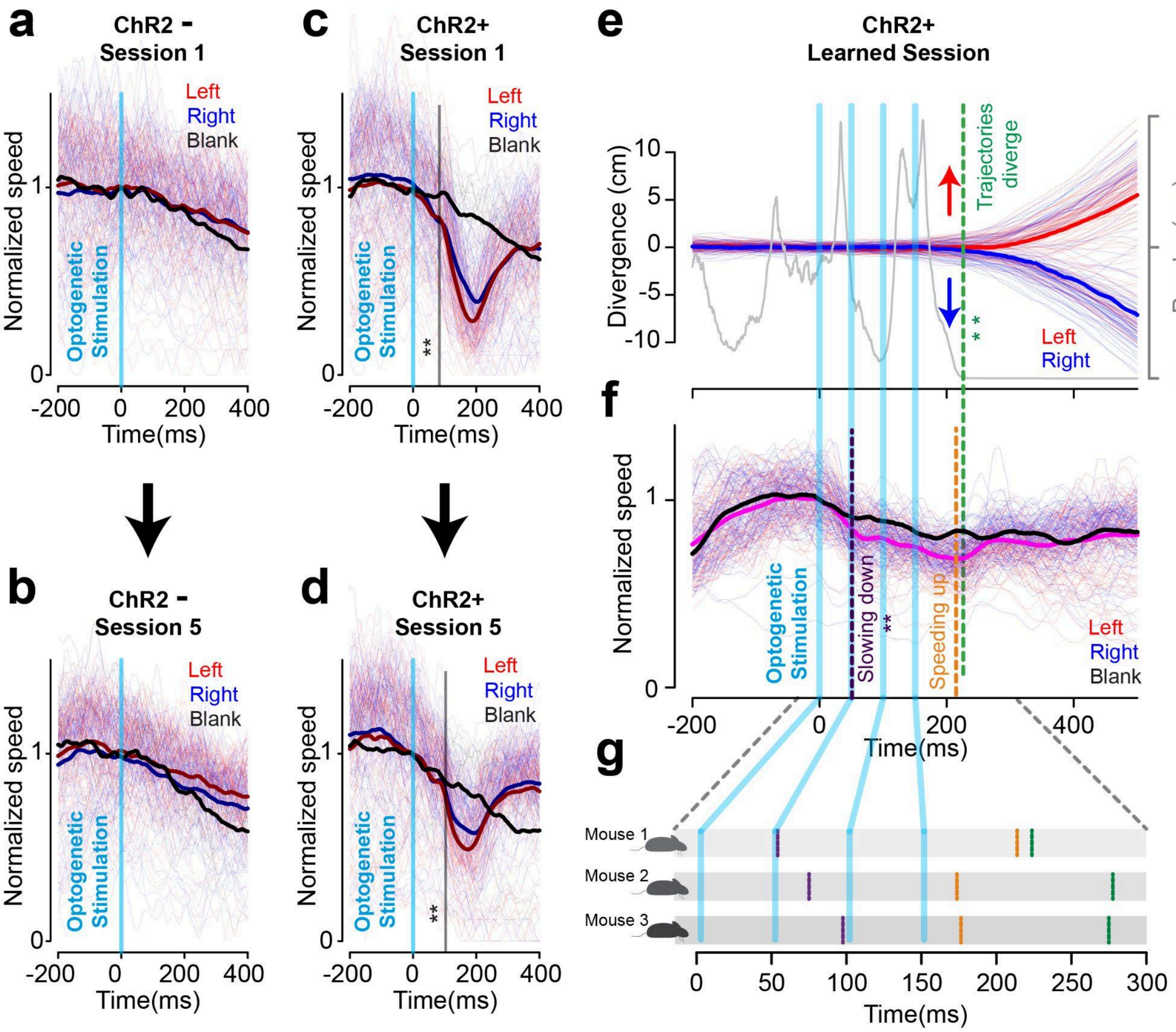


**Supplementary Figure 9. Sensory cortex stimulation induces motor effects. a)** Examples of normalized speed profiles (normalized to mean speed at stimulation time) during optogenetic and blank (no-stimulation) trials in a ChR2−(control) mouse navigating the large infinite Y-maze during session 1. Dark red, dark blue, and black lines indicate mean speed for left, right, and blank trials, respectively. **b)** Same as a), but for subsequent session 5 of the same ChR2− mouse. **c&d)** Same plots as in (a) and (b) but for a ChR2 + mouse. The vertical black line indicates the moment of speed of stimulated trials diverges from blank trials (statistically different, ** *P<0.01*). **e)** Aligned example trajectories from all intersections for a ChR2 + mouse during the learned session. Time zero is when the stimulation was presented. The gray line is the corresponding p-value for comparison of trajectory distributions for left (colored in red) and right (colored in blue) trials (Kolmogorov–Smirnov test) for every time point (1ms resolution). Vertical green line indicates when the two distribution are statistically different (** P<0.01). **f)** Normalized speed (to mean value at stimulation point) for both blank and stimulated trials for the same session where aligned trajectories are shown in (e). Pink line indicates the mean of optogenetic stimulation trials (left and right). Vertical purple lines represent the moment when speed of stimulation trials diverges from the speed for blank trials (Kolmogorov–Smirnov test, **P<0.01). Vertical orange lines (speeding up) indicate the moment when an animal's acceleration (derivative of speed) changes from negative to positive. **g)** Timeline for slowing down, speeding up and trajectory divergence (decision commitment) for all 3 ChR2+ mice in the large infinit Y maze.

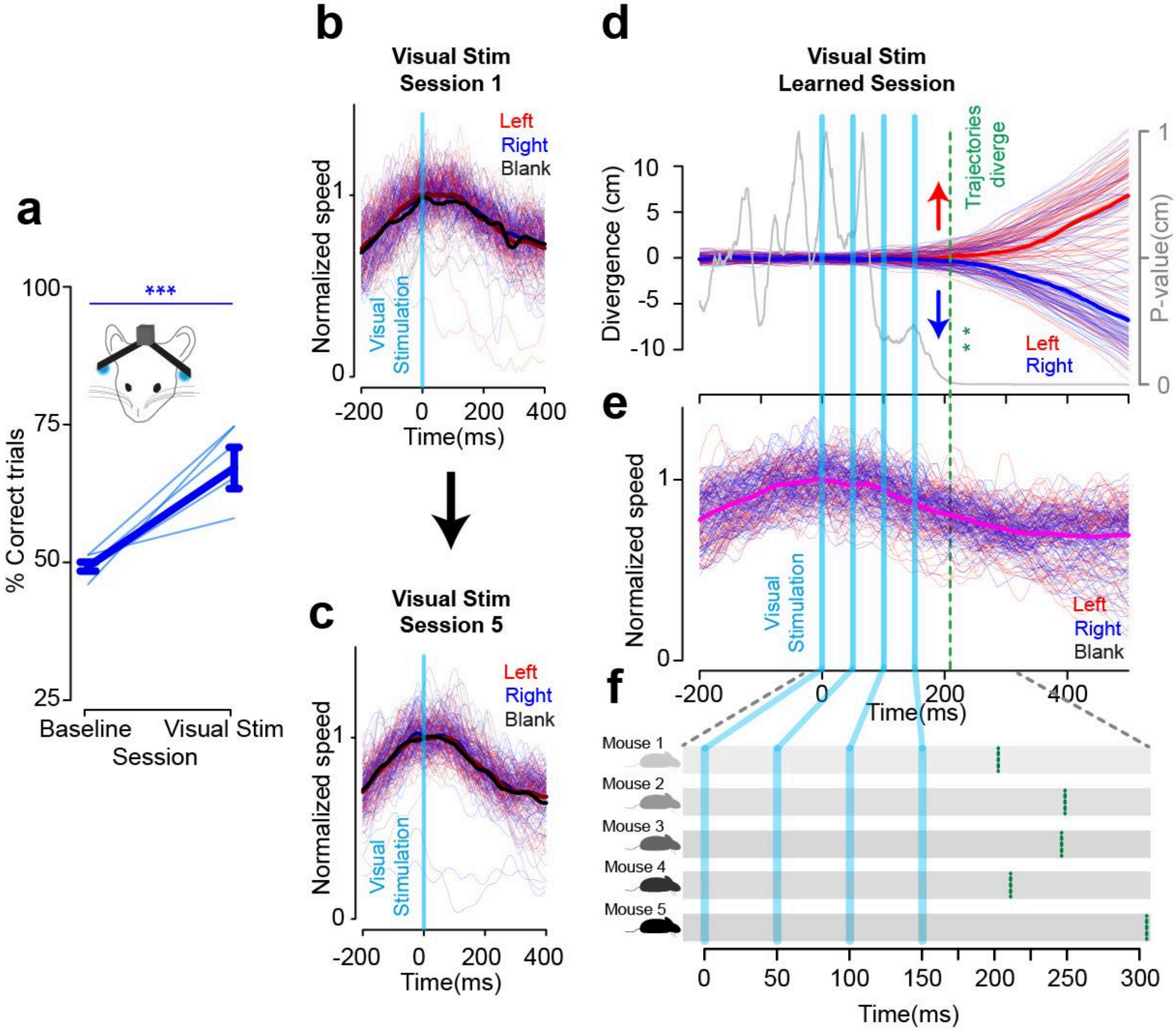


**Supplementary Figure 10. Closed-loop navigation control with visual stimulation. a)** Performance of mice in baseline (no stimulation) versus the last session to associate visual cues with the correct exit path (right led to exit from right arm and left LED to exit from left arm) at every intersection in large infinit Y maze (N=5 mice, one way repeated measure ANOVA, ***p<0.001). **b)** Example normalized speed traces (to mean value at stimulation point) during visual stimulation and blank (no visual stimulation) trials for a mouse in the first session navigating in the large infinit Y maze. Dark red, dark blue and black lines are the means for left, right and blank trials respectively. **c)** Speed profile same as shown in (b) but for session 5 of the same mouse. **d)** Example aligned trajectories from all intersections for a mouse during the learned session of visual association with exit path. Time zero is when the stimulation was presented. The gray line is the corresponding p-value for comparison of trajectory distributions for left (colored in red) and right (colored in blue) trials (Kolmogorov–Smirnov test) for every time point (1ms resolution). Vertical green line indicates when the two distribution are statistically different (** P<0.01). **e)** Normalized speed (to mean value at stimulation point) for stimulated trials for the same session where aligned trajectories are shown in (e). Pink line indicates the mean of visual stimulation trials (left and right). **f)** Timeline for trajectory divergence (decision commitment) for all 5 mice in the large infinit Y maze.

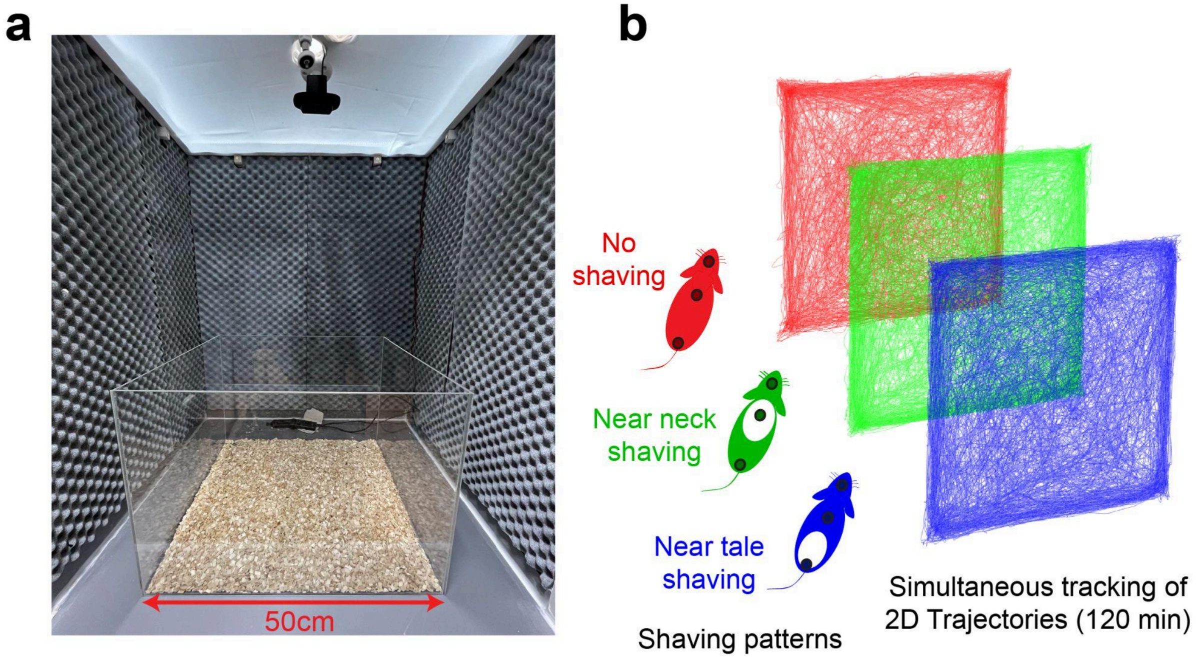


**Supplementary Figure 11. Experimental setup for closed-loop modulation of social behavior. a)** Arena for multi-animal setup. The transparent box is placed inside a wooden box covered with sound proof foam. **b)** Left) Shaving pattern applied to each cage of three mice to keep track of identity during DeepLabCut-Live tracking. The color code corresponds to the dominance hierarchy of mice within each cage as described in Figure 3. **b)** Example trajectories (120 minutes) simultaneous tracking of three mice using DeepLabCut-Live.

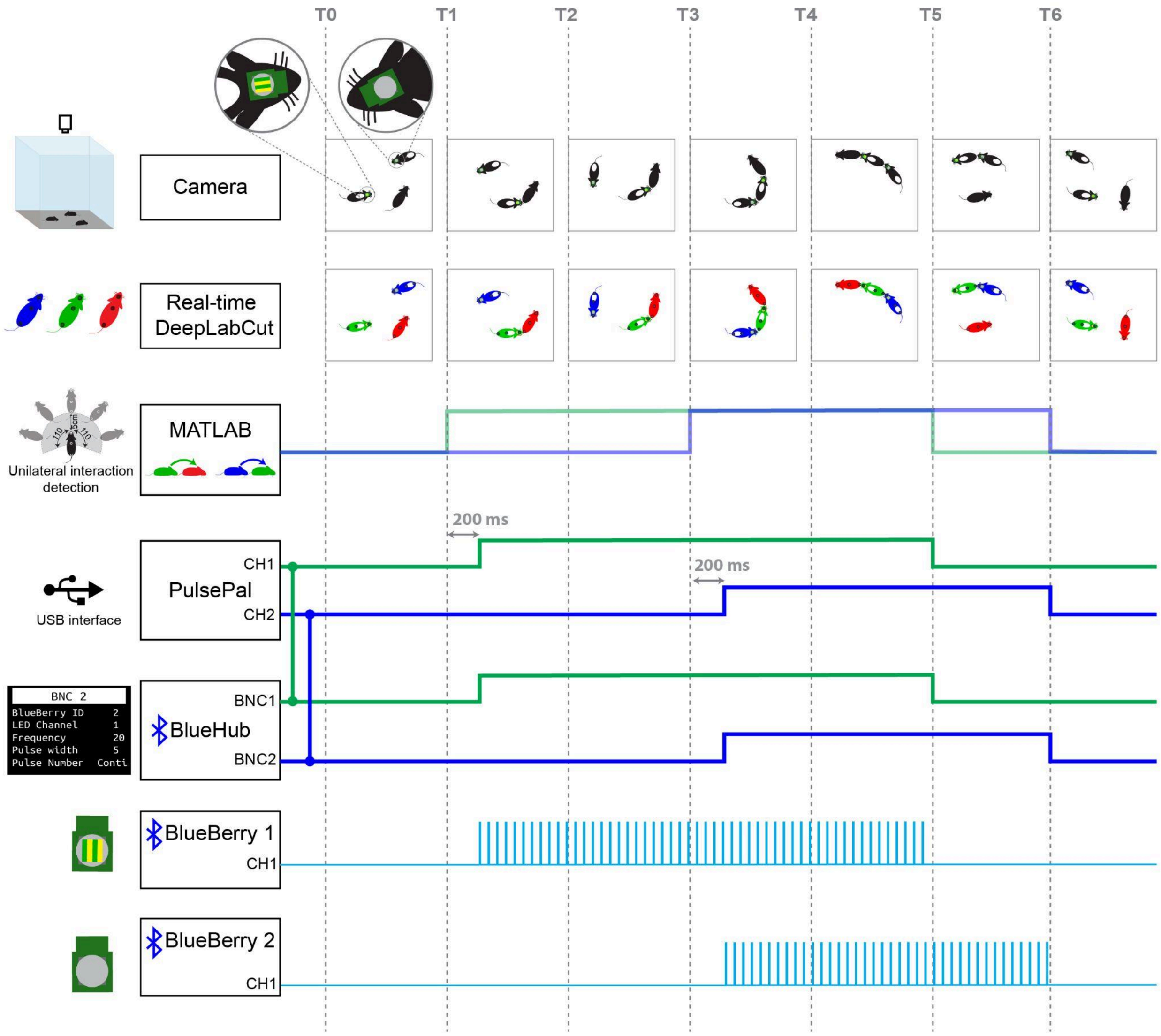


**Supplementary Figure 12. Pipeline for closed-loop modulation of social group dynamics.** Mice were first marked by individual shaving patterns and then placed in a 50 × 50 cm arena equipped with an overhead camera. For tracking accuracy and individual identification, one of the two BlueBerry devices (corresponding to the mouse with social rank 2, referred to as the “green mouse”) was marked with a piece of tape with 2 color striping patterns. DeepLabCut-Live tracked each mouse’s nose, body, and tail, while retaining their individual identities. These nine tracking points (three per mouse) were then processed in MATLAB to detect any unilateral interaction within specific pairs. Once such an interaction was initiated and maintained for over 200 ms, MATLAB sent a digital signal (5 V) via a PulsePal device. PulsePal’s two output channels were connected to BNC1 and BNC2 of the BlueHub, each pre-assigned to one of the two BlueBerry devices with stimulation parameters set at 20 Hz and 5 ms pulse duration. The BlueHub activated the assigned BlueBerry device(s) based on the incoming high signal and continued stimulation until detecting a return to a low digital level.

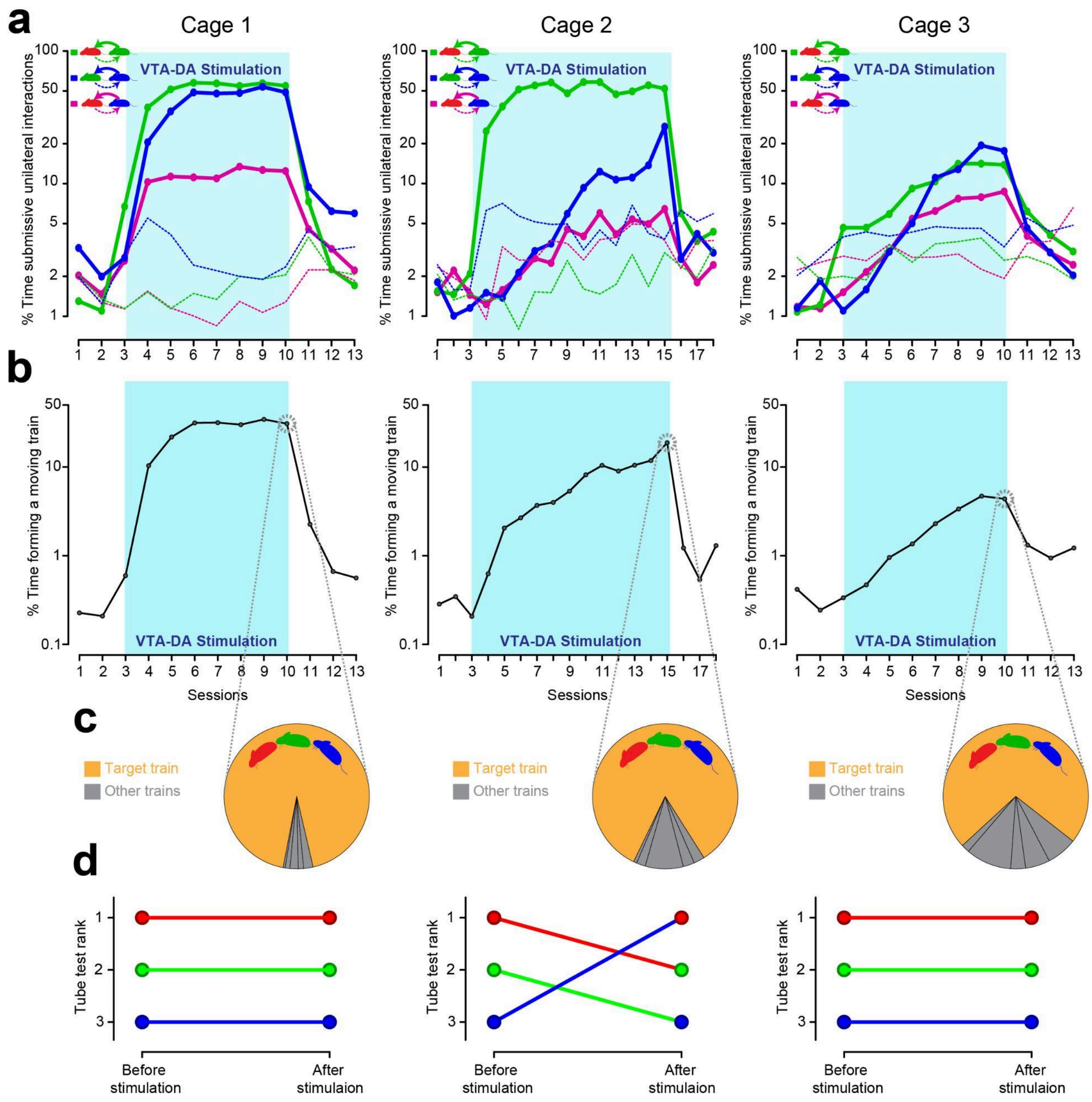


**Supplementary Figure 13. Detailed behavioral analysis for each cage during the social dynamics modulation experiment. a)** Percentage of unilateral dyadic interaction for each cage across pre stimulation, stimulation and post-stimulation sessions. Thick solid lines indicate unilateral interaction initiated by the submissive of each pair and the thin dashed lines are the unilateral interaction initiated by the dominant of each pair. The y-axis is scaled logarithmically. **b)** Train formation probability across pre-stimulation, stimulation and post-stimulation sessions for each cage. The y-axis is scaled logarithmically. **c)** Pie-chart representation of the distribution of all possible trains on the last day of VTA stimulation (with the target train shown in orange) for each cage. **d)** Social ranks for each cage determined through tube tests conducted both before day 1 and after the final day of stimulation.
